# Harnessing Plasmid Crosstalk in Transcription Factor Mediated Cell-Free Biosensors

**DOI:** 10.1101/2025.11.11.687754

**Authors:** Alexandra T. Patterson, Fernanda Piorino, Sasha Bronovitskiy, Diya Godavarti, Mark P. Styczynski

## Abstract

Plasmid crosstalk—unexpected changes to protein expression levels due to interactions between genetic cassettes in cell-free systems—complicates the creation of multi-plasmid cell-free tools. While the potential underlying mechanisms for crosstalk have been previously investigated, the practical impact of crosstalk on the implementation of cell-free genetic circuits has not been thoroughly examined. Here, we contextualize plasmid crosstalk in genetic circuits by examining its impact on the design and performance of multiple, diverse transcription factor biosensors. Guided by a deeper understanding, we harness crosstalk to enhance cell-free biosensor performance. Together, these findings demonstrate that plasmid crosstalk can serve as a tunable property in the development of cell-free genetic circuits.

## Introduction

Cell-free expression systems (CFE) have become a powerful platform for biosensing, with diverse applications spanning metabolic engineering, diagnostics, and environmental monitoring.^1–5^ Cell-free biosensors often harness the same sensing mechanisms as whole-cell biosensors but are easier to engineer and implement due to their shorter design-build-test-learn cycles.^1,2,6,7^ They are more compatible with field-deployment: they typically yield faster responses, they can be lyophilized and stored at room temperature with little to no loss of function, and their functions do not depend on cell viability or growth.^8–10^ The use of lysate-based cell-free biosensors, as opposed to their purified cell-free counterparts, is growing particularly rapidly, likely due to their lower cost, improved protein production, and higher compatibility with endogenous bacterial promoters.^2^

Transcription factor mediated biosensors, a major class of cell-free sensors, have been shown to efficiently detect a wide variety of small molecules.^11–14^ Generally, transcription-factor sensors are comprised of two parts: a regulator and a reporter. The regulator consists of an analyte-responsive transcription factor which modulates expression of the reporter (e.g., fluorescent or colorimetric protein) such that reporter production correlates with target analyte concentration. After identifying a regulator and reporter pair capable of detecting the analyte of interest, significant optimization is required to achieve desired biosensor characteristics, including fast response time, low signal without target (leak), and large dynamic range.^2^ Optimization is achieved though tuning the relative expression levels of reporter and regulator.^12,13,15,16^ In CFEs, the absence of a cell membrane allows this to be done with relative ease via adjusting concentration of regulator and reporter plasmid added into the reaction. To expedite this process further, the cassettes for the reporter and regulator are often coded for on separate plasmids, enabling orthogonal tuning of the two components.

However, expression of multiple plasmids in a single cell-free reaction may lead to off-target changes in protein levels, even with genetic cassettes that do not share regulatory interactions, a phenomenon known as plasmid-crosstalk. Our previous studies on crosstalk found that genetic cassettes will compete for transcriptional and translational resources in the CFE which may result in an increase or decrease in protein production. Promoter strength, promoter type, and plasmid dosage were shown to regulate the magnitude of crosstalk. Specifically, “strong” genetic cassettes (i.e., cassettes at high concentrations and/or cassettes with strong promoters) in the reaction generally monopolize transcriptional and translational resources, reducing expression from other “weaker” genetic cassettes (i.e., negative crosstalk). However, crosstalk may also cause an increase in expression via “nuclease distraction”. Briefly, the expression of another genetic cassettes can generate additional transcripts. These transcripts will sequester some of the nucleases, allowing more of the “original” transcripts to linger in the lysate and be subsequently translated (i.e., positive crosstalk). ^17^

CFEs involving regulatory interactions between genetic cassettes, such as transcription-factor sensors, are likely to have another layer of crosstalk complexity, as sensing capabilities are intertwined with plasmid concentration and expression. However, the extent of these factors on biosensor performance has yet to be investigated. A more detailed study on crosstalk would enable more informed design of genetic circuits, potentially reducing development time and improving biosensor functionality.

Here, we address this gap by investigating, characterizing, and harnessing plasmid crosstalk in the context of different transcription factor cell-free biosensors. We focus our initial investigation on three diverse, previously reported biosensors: a transcriptional repressor-based vitamin C sensor (repressor biosensor),^18^ an transcriptional activator-based vitamin B_12_ sensor (activator biosensor),^19^ and a dual-regulatory homocysteine sensor.^14^ We explore and compare the crosstalk landscape between different regulatory topologies, suggesting primary mechanisms of crosstalk’s effects on performance. Leveraging these insights, we propose and demonstrate the potential utility of crosstalk as an orthogonal biosensor tuning strategy. Through this novel approach, we demonstrate increases in dynamic range, fold-change, and achieve crosstalk-mediated biological computation. In total, these experiments demonstrate the impact, ubiquity, and potential utility of plasmid crosstalk in CFBs.

## Results and Discussion

### Characterization of Plasmid Crosstalk in a Repressor Biosensor

We first investigated crosstalk in the context of a previously reported vitamin C (ascorbate) biosensor.^18^ This biosensor harnesses transcriptional regulation by *E. coli*’s UlaR, which represses transcription from the endogenous P_UlaA_ promoter in the absence of ascorbate-6-phosphate.^20–22^ It includes two plasmids: the regulator, pUlaR, expressing UlaR from a T7 promoter, and the reporter, pUlaAsfGFP, expressing a reporter protein from P_UlaA_ (Figure 1A).

**Figure 1.**
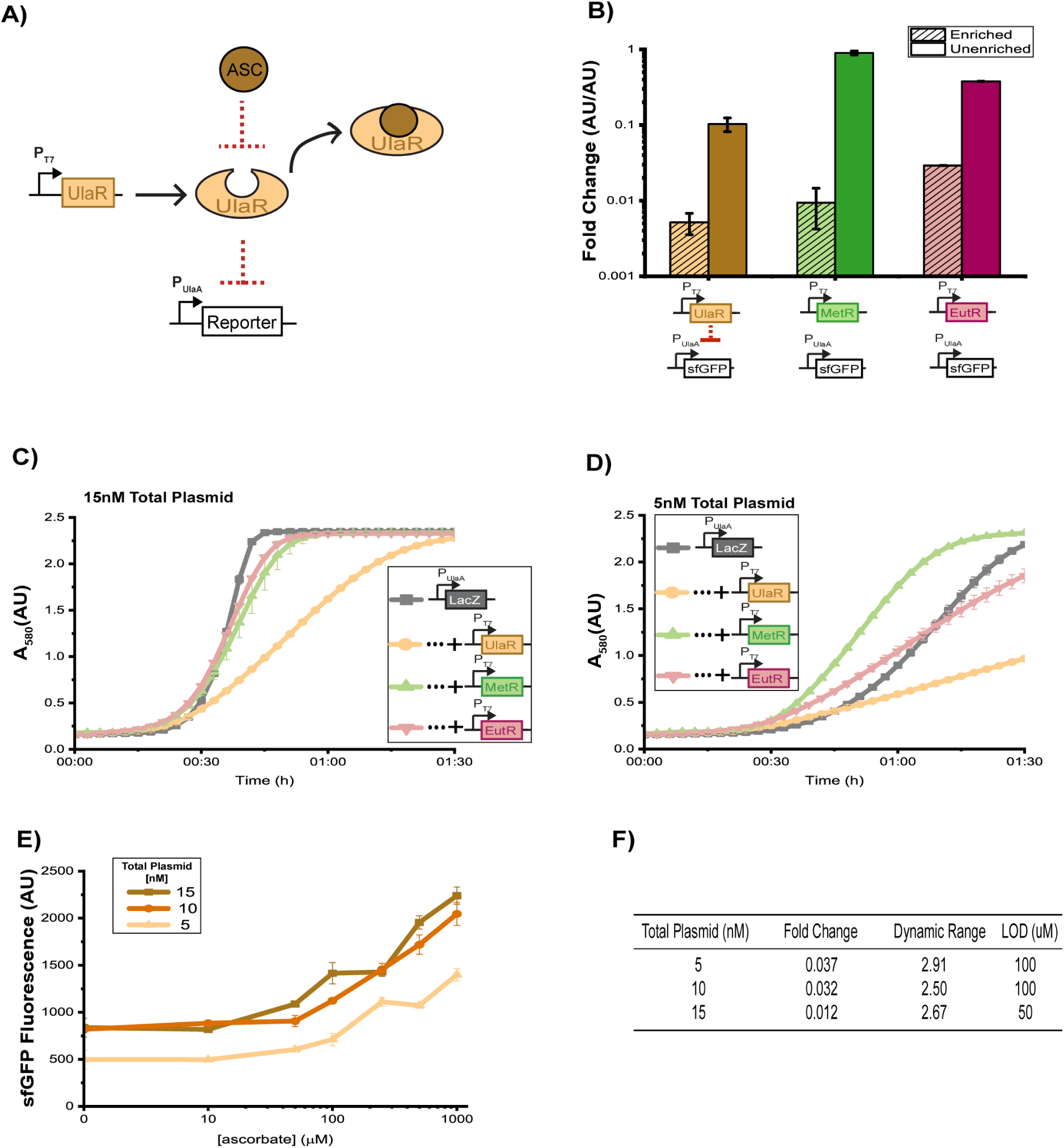
Investigation of plasmid crosstalk in a transcriptional repressor-based, cell-free vitamin C (ascorbate) biosensor. **(A)** Schematic of constructs used for the sensor. pUlaR expresses UlaR from a T7 promoter and P_UlaA_sfGFP expresses a reporter protein. When ascorbate is absent, UlaR represses expression from P_UlaA_. **(B)** Confounding effects of transcription factor expression and regulation on reporter output. Expression of UlaR, MetR, or EutR repressed pUlaAsfGFP, suggesting high-levels of negative plasmid crosstalk, as only UlaR should repress P_UlaA_-mediated transcription. In a lysate that is not enriched with T7 RNA polymerase, crosstalk is not as significant. MetR and EutR lead to different degrees of crosstalk, demonstrating that the identity of the protein being expressed matters. In all reactions, transcription factor plasmids were added at 5 nM, and P_UlaA_sfGFP was added at 10 nM. **(C)** Crosstalk in a vitamin C biosensor with a LacZ-based colorimetric reporter. Interactions between plasmids when transcription factor plasmids were added at 5 nM, and P_UlaA_LacZ was added at 10 nM. Crosstalk caused by expression of MetR and EutR is not as significant as with a sfGFP reporter. **(D)** Crosstalk in a vitamin C biosensor with a LacZ-based colorimetric reporter when transcription factor plasmids were added at 1.67 nM, and P_UlaA_LacZ was added at 3.33 nM.. Positive crosstalk is observed at all time points for expression of MetR and at early time points for UlaR and EutR. At later time points, UlaR-mediates strong repression. **(E)** Sensor response to a range of ascorbate levels at three total plasmid concentrations, maintaining a 2:1 ratio of Reporter:Regulator. The responses at 15 and 10 nM are qualitatively similar. At 5 nM, the sensor exhibits lower sfGFP expression in the absence of ascorbate, but also weaker fluorescent signal at high ascorbate levels. **(F)** Changes in the dynamic range and the detection limit of the vitamin C biosensor at different degrees of crosstalk based on Figure 1E. Decreasing total plasmid levels slightly reduces of magnitude of crosstalk, increases the detection limit, and improves the sensor’s dynamic range. In panels B, C, and D reactions were run in a T7 RNA polymerase-enriched lysate (unless otherwise stated). In panel E, a 50:50 mixture (by volume) of this T7 RNA polymerase-enriched lysate and a lysate enriched with phosphotransferase system proteins (UlaABC) was used. For sfGFP data, data were collected after 4 hours of incubation at 37°C. Background fluorescence was subtracted from all samples. For LacZ data, data were collected during incubation at 37°C. In all panels, error bars indicate the standard deviation of three technical replicates.

To begin delineating the effects of plasmid crosstalk in the UlaR biosensor, we analyzed the change in fluorescent signal from the reporter plasmid when either a cognate (pUlaR) or non-cognate transcription factor (pMetR, pEutR) was expressed (Figure 1B and S1, cross-hatched bars). Surprisingly, co-expression of both cognate and non-cognate transcription factors resulted in a significant reduction in fluorescence levels even though neither MetR nor EutR has been reported to mediate transcriptional repression of P_UlaA_. Thus, when MetR and EutR are expressed, this reduction in signal is likely attributable to negative crosstalk. In line with past reports on crosstalk, the T7 RNA polymerase driven expression of pUlaR, pMetR, and pEutR transcripts likely dominate translational resources, resulting in lower translation of the reporter plasmid.^17^

To investigate these phenomena further, we ran the same reaction in a lysate with basal (rather than induced) levels of T7 RNA polymerase. Here, we would expect to see a decrease in the extent of repression as fewer transcription factor transcripts are generated, freeing up expression resources for the reporter plasmid. In accordance with our hypothesis, co-expression in the non-enriched lysate greatly decreased the extent of repression. (Figure 1B and S1, solid bars). The reduction in fluorescent signal was most significant when UlaR was co-expressed in both the enriched and unenriched lysate, suggesting that UlaR can effectively repress PUlaA-mediated transcription. However, these results raise the question of a how effective of a repressor UlaR truly is, as the decrease in fluorescent signal upon addition of pUlaR seemingly arises predominantly from negative crosstalk. In addition, these data highlight the impact of the identity of the protein on crosstalk, as MetR and EutR seem to mediate very different degrees of crosstalk. This difference could potentially be attributed to codon usage, protein size, and amino acid profile, as these factors affect transcription and translation rates, but MetR and EutR have similar sizes (35.6 kD and 40.2 kD, respectively) and hydrophobicity (47.6% and 43.4%, respectively). Interestingly, while MetR caused stronger crosstalk than EutR in a T7 RNA polymerase-enriched lysate, the opposite was true in the unenriched lysate.

Noting the surprising difference in MetR and EutR repression, we wondered whether changing our reporter protein would shift the crosstalk landscape. We replaced the sfGFP gene in pUlaAsfGFP with LacZ, which encodes the β-galactosidase enzyme. This enzyme converts a yellow substrate (chlorophenol red-β-D-galactopyranoside, CPRG) to a purple product (chlorophenol red, CPR), causing an increase in absorbance at 580 nm. This approach generates a semi-quantitative readout with colorimetric outputs visible to the naked eye. Surprisingly, expression of MetR and EutR only slightly reduced absorbance values from the LacZ reporter, indicating a dramatic reduction in crosstalk-induced repression compared to the sfGFP reporter (Figure 1C). As expected, expression of UlaR induced the greatest repression, delaying terminal absorbance (around 2.25) by 45 minutes. Here, the observed repression is likely driven primarily by transcription-factor mediated repression rather than crosstalk, as evidenced by the absence of effects from MetR and EutR.

To further interrogate crosstalk with LacZ, we decreased total plasmid concentration to 5 nM while keeping the ratio of the reporter to the regulator constant at a 2:1 ratio (Figure 1D). Under these conditions, crosstalk again became a predominant influence, exhibiting both positive and negative regulation. At early time points, expression of all three transcription factors led to positive crosstalk, although to varying extents. This effect persisted for MetR until the terminal absorbance and is likely attributable to nuclease distraction, in which transcripts from the transcription factor gene predominantly occupy RNases, effectively increasing the half-life of leaky pUlaA transcripts. However, for EutR and UlaR, crosstalk effects eventually transition from positive to negative, as competition for translational resources may begin to outweigh the positive effects of nuclease distraction. In the case of UlaR, this crosstalk is combined with UlaR-mediated repression of PUlaA, which likely explains why absorbance levels are much lower than when EutR is expressed.

We next aimed to investigate the impact of crosstalk when sensing ascorbate. Specifically, we analyzed two key sensor parameters: dynamic range and limit of detection (LOD). In this work, we measured the dynamic range as the largest fold increase in reporter signal, and the limit of detection as the lowest analyte concentration that results in a reporter signal statistically different from the signal in the absence of analyte. Ideally, sensors should be able to detect analyte levels on the low end of the relevant concentration reference range with a dynamic range that allows for different levels within that range to be easily distinguished.

To evaluate key sensor metrics, we tested the sensor’s response to a wide range of ascorbate concentrations at different reporter and regulator concentrations (Figure 1E). Basal repression increased with plasmid concentration, likely due to an increased competition for resources, but did not result in substantial improvements in biosensor performance (Figure 1F). Based on our initial findings, a majority of the basal repression is likely attributable to crosstalk rather than transcription factor-mediated repression. Therefore, even upon ascorbate addition, system activation remains limited, as de-repression may be masked by negative crosstalk. Taken together, these results suggest that crosstalk has complex effects on sensor performance and can affect both activation and leak.

### Characterization of Plasmid Crosstalk in an Activator Biosensor

Having investigated crosstalk in the repressor-based vitamin C biosensor, we decided to examine crosstalk in the context of an activator mediated vitamin B_12_ biosensor, which harnesses EutR-mediated transcriptional activation in the presence of ethanolamine and vitamin B_12_ (here, we use the adenosylcobalamin—or AdoCbl—variant).^19,23^ In addition to the pEutR regulator, this sensor includes the reporter pEutSsfGFP (Figure 2A). This sensor was previously characterized in an extract with basal levels of T7 RNA polymerase, so we primarily used this lysate in the experiments described in Figure 2.

**Figure 2.**
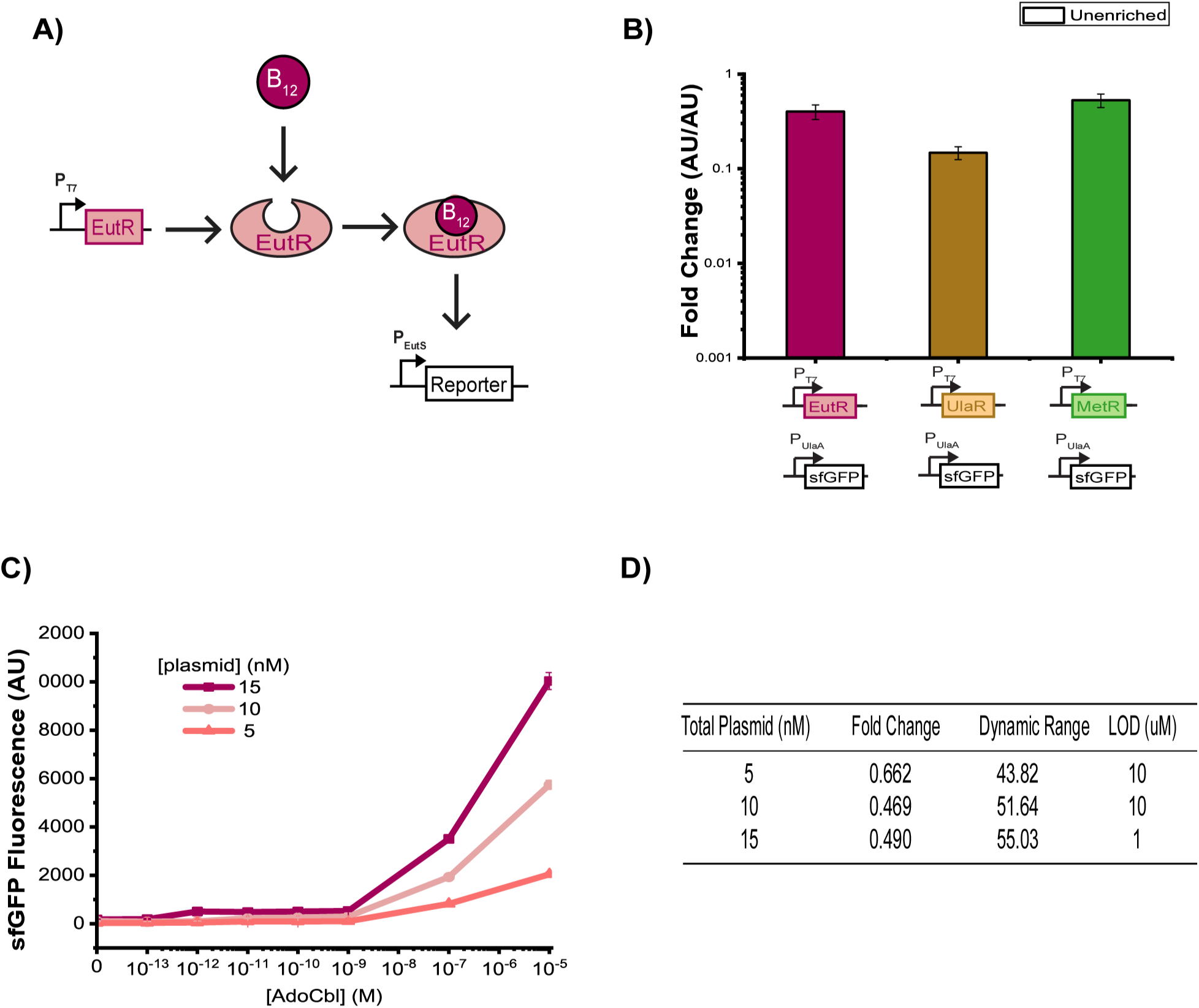
Investigation of plasmid crosstalk in a transcriptional activator-based, cell-free vitamin B_12_ biosensor. **(A)** Schematic of constructs used for the sensor. pEutR expresses EutR from a T7 promoter, and pEutSsfGFP expresses a reporter from P_EutS_. pEutR activates pEutS when vitamin B_12_ is present. **(B)** Confounding effects of transcription factor expression and regulation on reporter output. Repression from P_EutS_sfGFP when EutR, UlaR, or MetR is expressed from P_T7_, suggesting negative plasmid crosstalk as none of the transcription factors should repress P_EutS_-mediated transcription. In all reactions, 10 nM of pEutSsfGFP and 5nM of regulator were added. **(C)** Sensor response to a range of AdoCbl levels at three total plasmid concentrations, maintaining a 2:1 ratio of P_EutS_sfGFP:P_T7_EutR. The sensor’s response looks qualitatively similar at concentrations of AdoCbl below 1 nM, above which it exhibits significantly different dose responses. **(D)** Changes in the dynamic range and the detection limit of the vitamin B_12_ biosensor at different plasmid concentrations based on Figure 2C. The highest plasmid concentration tested (15 nM)—at which crosstalk is strongest—is correlated with the best dynamic range and the lower limit of detection In all panels, reactions were run in an unenriched lysate and data were collected after 4 hours of incubation at 37°C. 5 mM ethanolamine was added to all reactions. Background fluorescence was subtracted from all samples and error bars indicate the standard deviation of three technical replicates.

We began by analyzing the change in pEutSsfGFP signal when either a cognate (pEutR) or non-cognate transcription factor (pUlaR, pMetR) was expressed. In an activator topology, neither the cognate nor non-cognate transcription factor should result in any change to pEutSsfGFP expression, as regulation should only occur in the presence of AdoCbl. Addition of all three transcription factors resulted in a significant decrease in sfGFP expression, although to a much lesser extent when compared to the UlaR sensor (Figure 2B). Similar to the UlaR sensor, the three transcription-factors all repressed to different degrees, again highlighting the impact of protein identity on crosstalk.

We then analyzed biosensor performance in the context of crosstalk. As we did with the vitamin C biosensor, we tested a wide range of analyte levels at three total plasmid concentrations (Figure 2C). Interestingly, increased reporter and regulator plasmids improved biosensor performance, despite a higher magnitude of negative crosstalk (Figure 2D). This is in contrast to the Vitamin C sensor, where crosstalk needed to be minimized to improve functionality. In total, this highlights how crosstalk’s effects can vary across different regulatory networks.

### Characterization of Plasmid Crosstalk in a Dual-Regulatory Biosensor

We next sought to characterize crosstalk in a more complicated transcription-factor topology. Specifically, MetR has been shown to regulate the promoter P_GlyA_ in a homocysteine dependent manner.^14,24,25^ However, this sensor leverages both transcriptional repression (at low regulator and presumably low homocysteine concentrations) and transcriptional activation (at high regulator and high homocysteine concentrations).^26^ Previous attempts to use the P_GlyA_ promoter in a cell-free biosensor found it to have unacceptably high limits of detection and baseline leak, precluding its utility.^14^

To investigate this, we began by analyzing fold-activation of the sensor at different regulator and reporter plasmid concentrations. Specifically, the ratio of regulator to reporter was maintained at a 2:1 ratio, as the total plasmid concentration was adjusted (Figure 3A and S2). In agreement with previously reported data, we found the fold-activation to be suboptimal, maxing out at around 3. Notably, optimizing plasmid concentration yielded only modest changes in biosensor performance and sfGFP yield, suggesting an alternative bottleneck, such as crosstalk.

**Figure 3.**
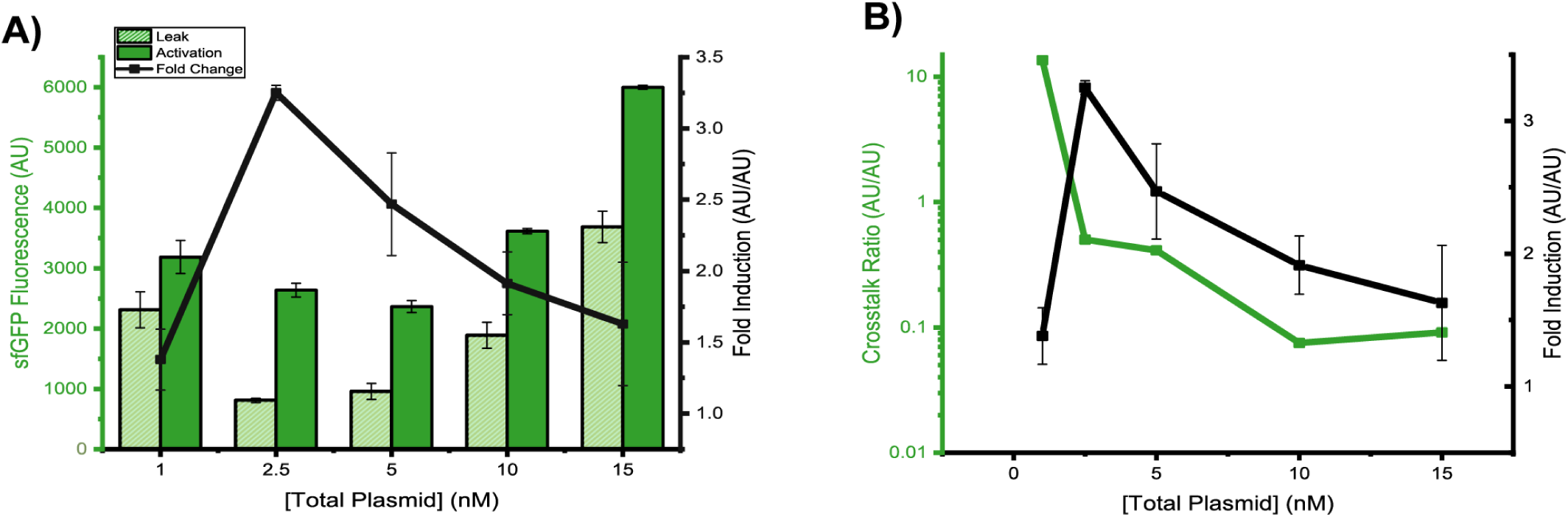
Investigation of plasmid crosstalk in a transcriptional dual-regulatory, cell-free homocysteine biosensor. **(A)** Leak (pGlyAsfGFP expression without homocysteine), activation (pGlyAsfGFP expression with homocysteine), and fold induction (activation / leak) at several plasmid concentrations. Fold induction peaks at 2.5nM total plasmid. **(B)** Correlation of crosstalk and leak at several plasmid concentrations. At 1 nM, pMetR causes positive crosstalk, resulting in decreased fold induction. From 2.5 nM to 15 nM, negative crosstalk increases and fold induction decreases. In all panels, data were collected in an unenriched lysate after 4 hours of incubation at 37°C. Reporter plasmid and pMetR were present at a 2:1 molar ratio. Error bars indicate the standard deviation of three technical replicates.

To quantitatively assess the degree of crosstalk, we divided the reporter output when both the reporter plasmid and the transcription factor plasmid are present by the reporter output generated by the reporter plasmid alone (crosstalk ratio) (Figure 3B and S2). Interestingly, at 1 nM plasmid, we observed strong positive regulation, such that the OFF state of the sensor (at 0 µM homocysteine) had a stronger fluorescent signal than the signal produced by pGlyAsfGFP alone. This results in an increase in baseline leak and a lower fold-change. As pGlyA is known to be act as a repressor at low concentrations, this regulation may be attributable to positive crosstalk. As plasmid concentrations increase from 2.5 to 15 nM there is a transition from positive to negative regulation, with the highest observed crosstalk levels occurring at the highest plasmid concentrations. However, given the dual-regulatory nature of MetR, it is a challenge to delineate whether the regulation can be attributable to crosstalk or transcription-factor regulation.

In all three topologies analyzed, there seems to be an optimal concentration of sensor plasmids that provides the ideal balance between crosstalk and regulator/reporter levels. Additionally. while the crosstalk landscape varied across the three biosensors tested, a few takeaways emerged. (1) Crosstalk is pervasive in transcription-factor sensors, spanning different regulatory topologies, multiple reporters, and across varied plasmid concentrations. (2) Transcription-factor crosstalk predominantly presents as negative crosstalk, reducing reporter expression. This phenomenon can likely be attributed to the weak, native reporter promoters typically used in transcription-factor sensors, as weak reporters have been previously shown to be highly susceptible to crosstalk.^17^ (3) Despite this, in some cases crosstalk led to improved biosensor phenotypes (e.g., decreased leak in the Vitamin B_12_ sensor). (4) However, changing plasmid dosage did not always affect biosensor performance in a predictable, linear fashion, highlighting the importance of carefully tuning and controlling for crosstalk between sensor plasmids.

It is worth noting that while qualitative trends are largely consistent across different lysate and plasmid DNA batches, we have noticed that the numeric magnitude of crosstalk may vary.^27^ In particular, the freshness of the purified plasmid DNA seems to matter considerably, with batches that have gone through multiple freeze-thaw batches or have been stored at 4°C for multiple weeks resulting in different degrees of crosstalk. Figure S3 reproduces the experiment of Figure 1B, but with freshly prepared plasmids. Even though the reactions were run with the same concentrations of plasmid and in the same lysate, fresh pUlaR and pEutR generated different degrees of crosstalk.

### Characterization of a Crosstalk-Mediated-Tuning Approach for Biosensor Optimization

Recognizing the potential of crosstalk to improve biosensor performance, we aimed to directly harness it as a tool in biosensor optimization. To minimize confounding effects of crosstalk on reporter and regulator function, we chose to generate crosstalk through a separate, non-regulatory plasmid (e.g., a plasmid expressing transcripts and/or proteins not directly involved in small molecule sensing or response). As a proof-of-concept, we began by tuning the functionality of a previously reported zinc biosensor.^11^ Briefly, *E. coli*’s ZntR activates transcription from the endogenous P_ZntA_ promoter in the presence of zinc. It includes two plasmids: the regulator, pZntR, expressing ZntR from a T7 promoter, and the reporter, pZntALacZ, expressing LacZ from P_ZntA_ (Figure 4A).^28^ All tuning experiments were conducted in an optimized base system, where reporter and regulator concentrations were optimized to yield strong activation and low leak at 5 µM zinc with a 40 minute incubation time, demonstrating that crosstalk-mediated tuning (CMT) can operate as an additional optimization tool in biosensor development.

**Figure 4.**
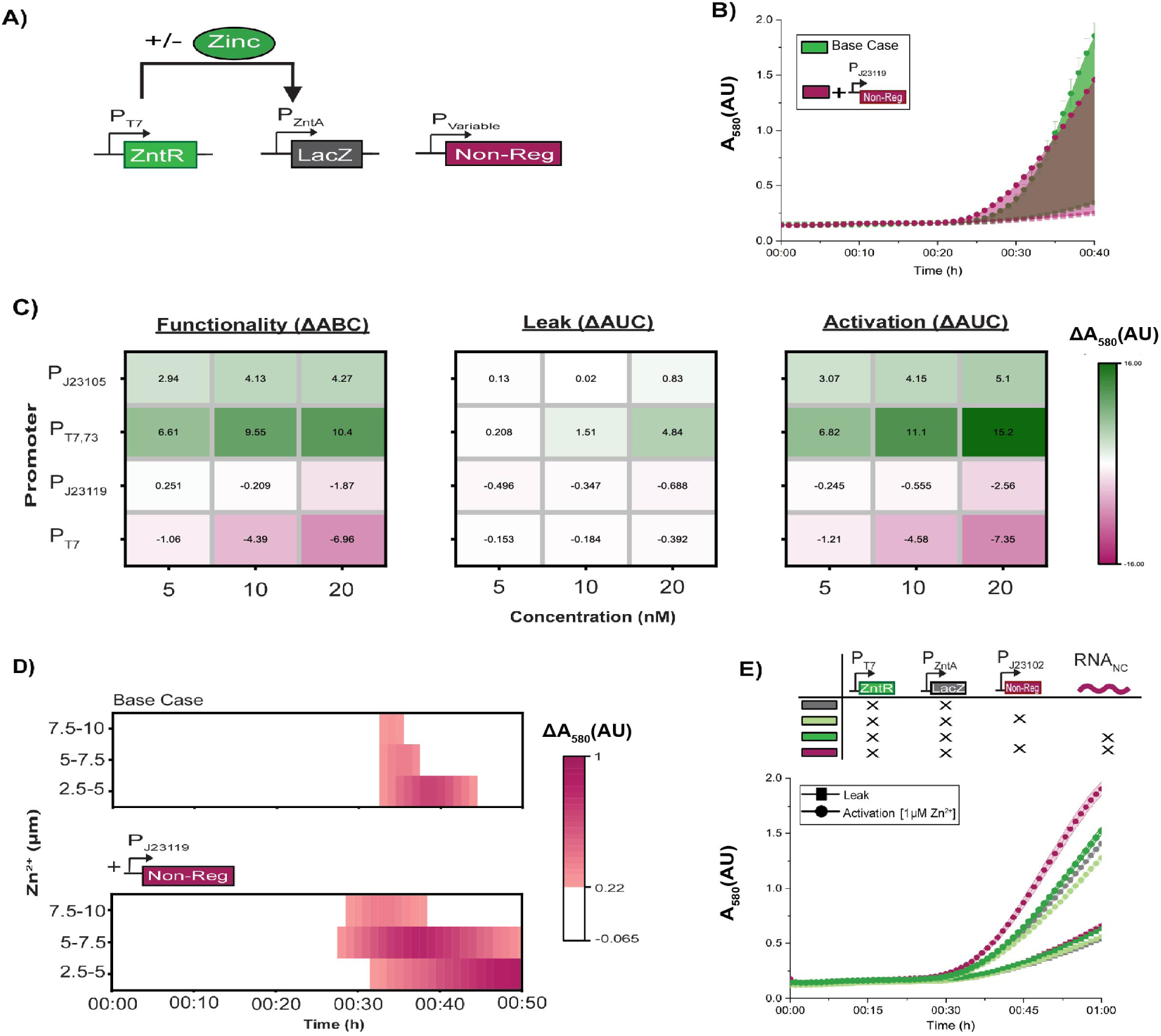
Investigation of crosstalk-mediated-tuning as an orthogonal optimization strategy in transcription-factor biosensor development. **(A)** Schematic of constructs used for the sensor. pZntR expresses ZntR from a T7 promoter and P_ZntA_LacZ expresses a colorimetric reporter protein. When zinc is present, ZntR activates expression from P_ZntA_. **(B)** Activation and leak upon addition of a non-regulatory sfGFP plasmid driven by P_J23119_. At early-time points pJ23119-NR induced positive crosstalk in the activation condition. At later time points, pJ23119-NR induced negative crosstalk in both the activation and leak condition. **(C)** Comparison of overall functionality, leak, and activation when the non-regulatory plasmid is expressed at different concentrations and under different strength promoters. Crosstalk-mediated-tuning appears to be highly correlated with both plasmid strength and concentration. Generally, weak promoters induce positive crosstalk and strong promoters induce negative crosstalk. **(D)** Effect of 10nM pJ23119-NR on the ability to distinguish between different zinc concentrations. In accordance with previously published results, zinc concentrations were considered indistinguishable if the difference in A_580_ readings between two concentrations was less than 0.22. Indistinguishable A_580_ readings are denoted in white.^11^ Times highlighted in red indicate visual detection, as the absorbance difference between the two zinc concentrations on the y-axis is above the 0.22 threshold. Darker colors represent a larger difference in A_580_, enabling easier interpretation. The addition of pJ23119-NR reduces incubation time, increases the time in which colors can be distinguished, and increases the absorbance difference between different zinc concentrations. **(E)** At low zinc concentrations (1 nM), crosstalk can be used to increase signal. Here, crosstalk was achieved by adding pJ23102sfGFP at 0.5 nM and 250 nM of non-coding RNA. In all panels, the reporter plasmid was added at 0.5 nM and the regulator plasmid was added at 5nM. The reactions were performed in a non-enriched lysate to increase competition for the T7 RNA polymerase. Unless otherwise noted, zinc was added at 5 µM. Data were collected during incubation at 37 °C. Error bars indicate the standard deviation of three technical replicates.

We began assessing CMT by measuring activation and leak upon addition of a non-regulatory plasmid expressing sfGFP under a strong, constitutive σ^70^ promoter (pJ23119-NR) (Figure 4B). At early-time points pJ23119-NR induced positive crosstalk in the activation condition but did not affect leak, ultimately resulting in a faster time to detection. Here, pJ23119-NR may amplify initial signal from the zinc activated pZntA promoter via nuclease distraction. In line with previous findings on crosstalk, the non-activated pZntA promoter (i.e., leak) may not produce the minimum number of transcripts required to induce positive crosstalk, resulting in no observed change in response. At later time points, crosstalk appears to transition from positive to negative possibly due to competition for ribosomes as the non-regulatory transcripts begin to accumulate.

To explore this phenomenon further, we analyzed the effects of non-regulatory promoter strength and plasmid concentration on CMT. Specifically, we analyzed CMT at three different plasmid concentration—5, 10, and 20nM—with four different promoters—strong σ70 (P_J23119_), weak σ70 (P_J23105_), strong T7 (P_T7_), and weak T7 (P_T7,73_) (Figure S4, S5). To aid in analysis, we adapted a previously published strategy to one-dimensionalize temporal LacZ biosensor response.^29^ Briefly, the area under the curve (AUC) was calculated for the leak and activation curves. CMT effects on leak and activation were quantified as ΔAUC=AUC_NonRegulatory_-AUC_Base_. To quantify the total effect of crosstalk on biosensor functionality, a similar strategy was employed; the area between the activation and leak curves (ABC) under both conditions (leak and crosstalk) was calculated, with ΔABC=ABC_NonRegulatory_-ABC_BaseCase_. For both ΔAUC and ΔABC a value above 1 corresponds with positive crosstalk and a value below 1 corresponds with negative crosstalk (Figure S4, Figure S5).

The different plasmid concentrations and promoters induced a wide variety of responses in the ΔABCs, both improving and impairing biosensor performance (Figure 4C). All combinations of concentrations and promoters tested resulted in some degree of crosstalk, although the quantitative extent of crosstalk varied greatly. Generally, strong promoters (P_T7_ and P_J23119_) led to negative crosstalk and weak promoters (P_T7,73_ and P_J23105_) led to positive crosstalk. This observation is consistent with our understanding of crosstalk, as stronger promoters monopolize transcriptional and translational resources, outweighing the positive effects of nuclease distraction achieved with weaker promoters. Furthermore, higher plasmid concentrations generally increased the extent of crosstalk, either in the positive or negative direction, depending on promoter strength. Here, higher concentrations of plasmids can achieve higher transcriptional rates, which can increase the extent of resource competition or nuclease distraction. Similarly, T7 promoters induced larger effects of CMT, as they have higher transcriptional rates when compared to their sigma70 counterparts. ^17^

To gain greater insights into the mechanisms of CMT, we also analyzed individual differences in the ΔAUC for activation and leak (Figure 4C). The effects of CMT are more apparent for activation compared to leak, which may be attributable to regulator and reporter concentrations being initially optimized for minimal leak, dampening the potential effects of CMT. Furthermore, both activation and leak follow similar trends to the ΔABCs. Therefore, large increases in activation are primarily accompanied by increases in leak, neutralizing the effect of positive CMT on system performance. Conversely, large decreases in activation are primarily accompanied by decreases in leak, slightly buffering any negative effects of CMT on system performance.

A prominent exception to these trends is observed when using P_J23119_, as increasing concentrations of plasmid led to a shift from positive to negative ΔABC. Interestingly, all concentrations of P_J23119_ plasmids showed positive crosstalk at early time points, which may be driving this unique CMT trend (Figure S6). Specifically, at 5 nM pJ23119-NR ΔAUC values for both the leak and activation condition decrease. However, activation is affected to a lesser extent as it is buffered by early positive crosstalk, resulting in a modest increase in the overall system performance as indicated by the increased ΔABC. As the concentration of pJ23119-NR is further increased, early positive crosstalk is no longer able to buffer against the larger decreases in end-point absorbance, leading to a reduction in overall system performance. We do note that early positive crosstalk can also be observed, albeit to a lesser extent, with low concentration of non-regulatory plasmid driven by P_T7,_ suggesting that this phenomenon is not unique to P_J23119,_ and may be a function simply of transcriptional strength.

### Applications of Crosstalk-Mediated-Tuning in Biosensor Development

After characterizing the landscape of CMT, we aimed to apply it to solve challenges in biosensor development which are not easily addressed using conventional reporter and regulator tuning. For example, to achieve semi-quantification of biomarkers at the point-of-care, it can be critical to maximize the color change between different biomarker concentrations and maximize the interval of time in which different concentrations can be visually discerned. In practice, these approaches make semi-quantification significantly more robust at the point-of-care, by minimizing misinterpretations from subjective color interpretation and user-variability in reaction timing. However, this is not always a straightforward task. Specifically, previous attempts to modulate the zinc sensor found that decreasing reporter plasmid concentration could maximize the time interval of which a dose-dependent zinc response is interpretable, but significantly increased the assay time.^11^ Therefore, it would be beneficial if CMT could be used to maximize color differences across zinc concentrations without increasing incubation time.

Noting the unique temporal dynamics of CMT when the non-regulatory plasmid was driven by P_J23119_, we hypothesized that positive CMT at early time-points could reduce incubation time, while negative CMT, which occurred only at later time points, could slow the reaction, such that color distinction is visible for a longer period of time. To test this theory, we added different concentrations of zinc with and without pJ23119-NR and compared visual discernment between zinc concentrations. In accordance with our hypothesis, the addition of non-regulatory plasmid not only greatly extended the time interval in which dose-dependence could be seen, but also decreased the time to detection (Figure 4D). This approach also increased the absorbance change between zinc concentrations, making visual interpretation even more robust.

Furthermore, we postulated that positive CMT could be strategically applied to maximize fold-change when detecting low levels of analyte, another common optimization challenge in biosensor development. To achieve this, we used a two-pronged CMT system to selectively increase activation without increasing leak. The first prong induces slight negative crosstalk via a non-regulatory plasmid under the control of a medium strength σ70 promoter (P_J23102_). Low concentration of this plasmid alone slightly decreased signal from the activation condition without a visible change in leak (Figure 4E, light green lines). The second prong induces low levels of positive crosstalk via purified non-coding RNA (i.e., RNA with no ribosomal binding site), enabling nuclease distraction without concerns for polymerase or ribosome competition. As expected, low concentrations of non-coding RNA alone slightly increased signal from both the activation and leak condition (Figure 4E, dark green lines). However, when both the non-coding RNA and P_J23102_ plasmid were added, a significant increase in activation occurred, without further increases in leak (Figure 4E, pink lines). Here, we hypothesize that the combination of non-coding RNA and zinc works to effectively increase the strength of P_ZntA_, such that it is stronger than P_J23102_. This shifts the crosstalk regime by enabling activated P_ZntA_ to overcome resource competition, such that presence of P_J23102_ induces positive CMT. Conversely, non-activated P_ZntA_ is much weaker than P_J23102_, such that CMT effects are negligible. In total, these two approaches show the potential of crosstalk to be harnessed as an orthogonal optimization tool in biosensor development.

### Towards a Crosstalk-Mediated Inverter

After demonstrating the utility of CMT in biosensor optimization, we aimed to induce greater changes in functionality by developing a crosstalk-mediated inverter (i.e., a single-input-single-output NOT logic gate). As a proof-of-concept, we aimed to invert signal from the previously published Zur repressor.^30^ Briefly, *E. coli*’s Zur represses transcription via binding to the Zur operon, blocking the polymerase from the promoter. The system includes two plasmids: the regulator, pZur, expressing Zur from a T7 promoter, and the reporter, pZurO, which can be regulated by any promoter when placed upstream from the ZurO site (Figure 5A).^31^ To modify this system into a crosstalk-mediated inverter, the reporter was moved to a third constitutively expressed plasmid (pCMI) and the zinc regulated plasmid, pZurO, was modified to express a non-regulatory protein (pZurO-NR) (Figure 5B). Given crosstalk’s ability to redirect transcriptional and translational resource, we postulated that high levels of transcription from pZurO-NR would monopolize transcriptional and translational resources, indirectly repressing the pCMI via plasmid crosstalk. However, in the presence of zinc, pZurO-NR should be repressed via ZuR, releasing resources back to pCMI, thus inverting the signal upon addition of zinc. To induce inversion, a 10:1 ratio was maintained between pZurO-NR and pCMI, in an attempt to pull as many resources as possible away from the low concentration pCMI plasmid. Excitingly, we found a strong inversion of signal when pCMI was expressed under P_T7_ (Figure 5C).

**Figure 5.**
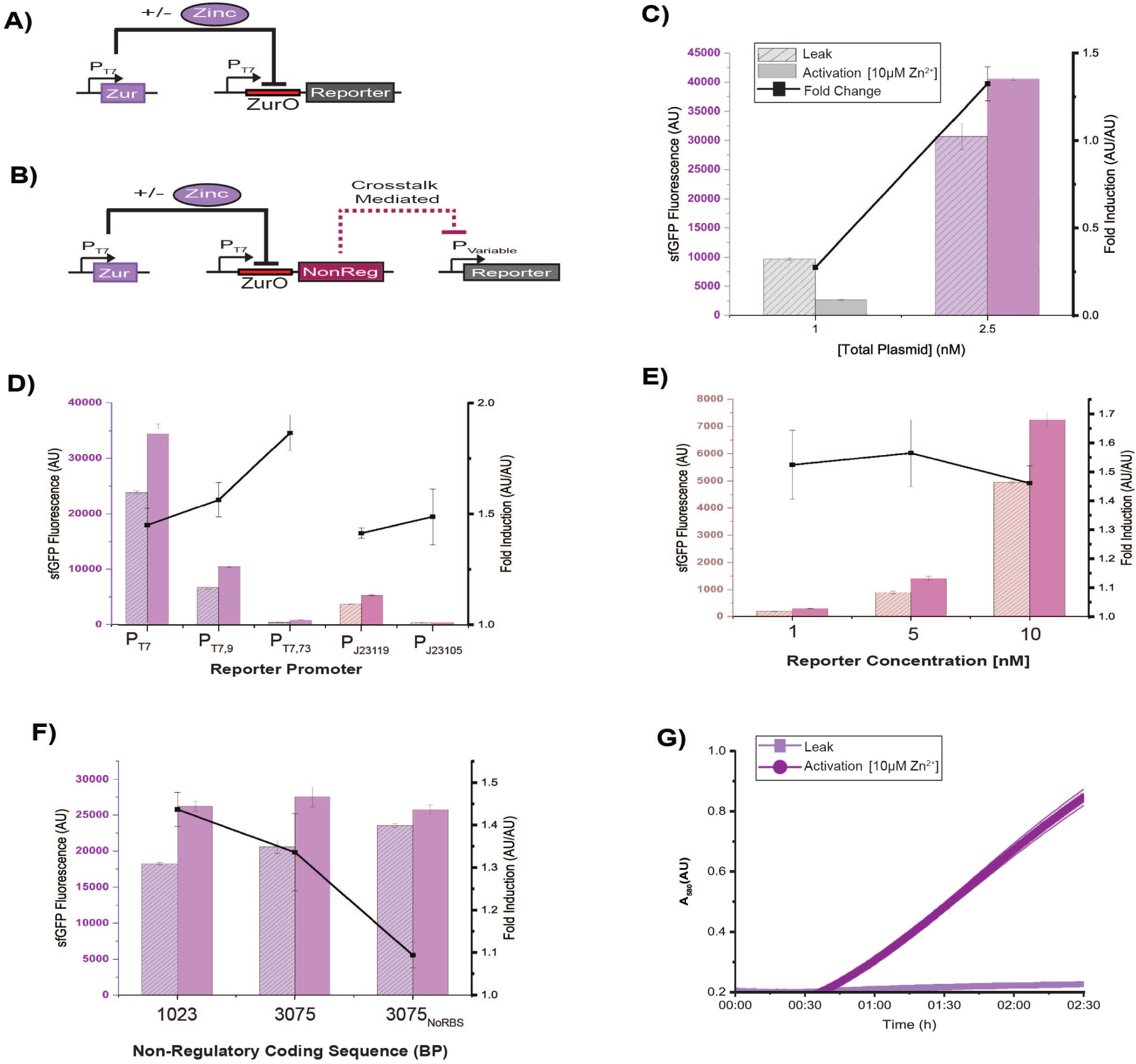
Investigation of crosstalk-mediated-tuning approach to invert signal from a repressor-based transcription-factor cell-free biosensor. **(A)** Schematic of constructs used in the base case repressor system. pZur is expressed from a T7 promoter and represses the reporter via binding to ZurO in the presence of zinc. **(B)** Schematic of constructs used in the crosstalk-mediated inverter. pZur is expressed from a T7 promoter and represses the expression of a non-regulatory in the presence of zinc. When repressed, resources are shifted away from the non-regulatory protein and towards the reporter plasmid, giving the appearance of activation in the presence of zinc. **(C)** Comparison of biosensor response between the base system and the inverter. The crosstalk-mediated inverter design is capable of inverting signal upon addition of zinc. **(D)** Comparison of inversion when the reporter is expressed under different strength promoters. Weak T7 promoters result in the highest fold induction when zinc is added. **(E)** Comparison of inversion when the reporter is added to the reaction at varying concentrations. Concentration does not appear to have strong effects on fold induction. **(F)** Comparison of inversion with different non-regulatory components. Translation, specifically translation initiation, appears to be critical for strong inversion. **(G)** Inversion with a LacZ-based colorimetric reporter. sfGFP was replaced with LacZ in the reporter plasmid. Crosstalk-mediated inversion was functional in a colorimetric sensor when reporter plasmid concentration was low (0.0001 nM). Unless otherwise noted, the reporter plasmid was expressed under P_T7_ and added to the reaction at 1 nM. In all panels, pT7Zur was added at 1nM and zinc was added at 10 µM. For panels A-F, data were collected at 3 hours after incubation at 37°C. For panel G, data were collected during incubation at 37 °C. Error bars indicate the standard deviation of three technical replicates.

To further interrogate the mechanisms of inversion, we altered promoter strength and plasmid concentration of pCMI. In total, all combinations of promoters and concentrations resulted in some degree of inversion, although the extent of inversion was variable. Weaker T7 promoters resulted in decreased leak (i.e., uninduced expression) from pCMI and slightly increased fold-repression (Figure 5D, purple bars). This is likely attributable to pZurO-NR being at both a high concentration and under strong P_T7_, pulling away transcriptional and translational resources from the weaker P_T7_ promoters. This reduces the leak of pCMI through negative crosstalk, and in-turn may facilitate a greater fold-change when de-repressed. As σ70 promoters are weaker than their T7 counterparts, lower expression levels are expected (Figure 5D, pink bars). However, strength of the σ70 promoters showed a less clear correlation with fold-repression. Similarly, increases in pCMI-P_J23119_ concentration resulted in higher uninduced expression but did not cause significant changes in fold-repression (Figure 5E). Here, higher pCMI concentrations likely lead to reallocation of translational resources towards pCMI, increasing the uninduced expression.

Next, we investigated how the transcriptional and translational dynamics of pZurO-NR can affect inversion. We began by changing the size of the pZurO-NR expressed protein. We hypothesized that larger proteins would dominate more translational resources, leading to lower uninduced expression and a higher fold-repression. Surprisingly, a 3x change in protein size led to a modest decrease in fold-repression and a small but significant increase in leak (Figure 5F). This counterintuitive result may suggest that translation initiation monopolizes more resources when compared to elongation. Therefore, having a larger number of shorter transcripts, causes greater resource coemption due to the increased number of RBSs. This finding is in line with whole-cell studies which have shown that initiation is the rate limiting step in translation.^32,33^ Furthermore, to interrogate the impact of transcriptional vs translational competition in inversion, we removed the RBS on the pZurO-NR, effectively isolating the effects of transcriptional competition. Removal of the RBS significantly increased leak, resulting in a reduced fold-repression (Figure 5F). These results suggest that transcriptional competition is not significant in this system.

Last, we wanted to explore the modularity of this approach by switching to a colorimetric reporter. To accomplish this, sfGFP was replaced with LacZ in pCMI. Unexpectedly, at high concentrations of pCMI-LacZ we saw faster color-change when zinc was added when compared to the base case (Figure S7). However, when pCMI-LacZ concentration was significantly reduced, clear inversion could be seen. These results are in-line with Figure 1, where crosstalk effects were less significant when using a LacZ reporter compared to a sfGFP reporter. We do note that by reducing the pCMI-LacZ concentration, the reaction was significantly slowed but had virtually no leak.

## Conclusions

Here, we have demonstrated the impact of plasmid crosstalk on the performance of three transcription factor-based cell-free biosensors. We found that crosstalk is highly prevalent in transcription-factor mediated CFEs and can greatly affect apparent performance. We showed that plasmid-driven expression of transcription factors causes a substantial decrease in reporter protein output. This negative crosstalk can be conflated with transcription factor-mediated regulation; it masks the true regulatory effect of the protein and—in the absence of any regulatory interactions— may give the appearance of regulation. Varying plasmid concentrations, reporter protein, and transcription-factor architecture changed the degree of crosstalk, and also led to changes in the dynamic range and limit of detection of the sensors.

We have also shown that the degree of crosstalk is not solely correlated with the transcriptional and translational elements of a genetic cassette. While the impact of competition of transcriptional and translational resources provides an initial guide for the design of different genetic cassettes, it does not explain some of our more nuanced results, such as the fact that expression of different proteins from the same promoter and RBS generated very different degrees of crosstalk. Taken together, these results highlight the importance of individually assessing crosstalk effects for a particular system, as these effects do not seem to be directly translatable across different systems even if the protein expression machinery is the same.

Enabled by a deeper understanding of crosstalk’s impacts on transcription factor biosensors, we were able to intuitively harness crosstalk as an orthogonal tuning strategy in CFEs. Through this strategy, we unlocked diverse and advantageous biosensor functionality that can be challenging to achieve using conventional tuning approaches, including increasing the signal-to-noise ratio and decreasing incubation time. We were able to further apply crosstalk for complex biological computation by demonstrating its ability to function as an inverter. To the best of our knowledge, this is the first demonstration of crosstalk being directly harnessed as a tool in CFE.

While this work focused on characterizing and modulating crosstalk in transcription-factor biosensors, our results suggest that crosstalk is an important design consideration in virtually all CFE applications requiring the expression of multiple proteins. At minimum, these data demonstrate the importance of considering and controlling for crosstalk when tuning multi-gene expression in CFE. Additionally, the orthogonal tuning approach developed here could be advantageous in many CFE system’s requiring tight control of multi-gene expression. For example, in CFE biomanufacturing it is often necessary to express multiple enzymes at precise levels over the course of a CFE reaction.^34^ Here, crosstalk-based tuning could be coupled with transcription-factor sensors to respond and regulate enzyme production in response to changes in metabolite levels. Similarly, CFE crosstalk tuning could be applied in genetic circuit prototyping,^35,36^ protein prototyping,^37^ or synthetic cell development,^38,39^

However, for crosstalk to be efficiently utilized as an orthogonal tuning strategy, further work must be done to standardize and model CFEs. Computational models that directly account for crosstalk would be of great utility and could allow us to more accurately predict interactions between genetic cassettes and their effect on protein outputs. Recent work has made progress towards a mathematical model of crosstalk.^40^ However, challenges in lysate and plasmid variability and unwieldy parameter identification still preclude a quantitively predictive model. ^27,41,42^ Furthermore, crosstalk is a complex phenomenon, and additional experimental efforts are still needed to elucidate all its underlying mechanisms. Nonetheless, the results reported here are already important, as they provide a basis for incorporating crosstalk into the design and analysis of cell-free platforms

In total, these results show the pervasive, underexplored nature of CFE in a diverse set of regulatory circuits. Our results suggest that crosstalk is likely an inevitable feature of multi-plasmid expression in CFE. However, through intuitive design, guided by an in-depth characterization, we showed crosstalk can not only be overcome but harnessed as a tool to achieve finely-tuned multi-protein expression. As CFE applications move towards increasingly complex applications, and in-turn require more robust and predictable multi-protein expression, there will be a growing need to fully elucidate, control for, and in some cases harness plasmid crosstalk.

## Methods

### Materials

T4 DNA ligase, T5 exonuclease, Taq ligase, Phusion DNA polymerase, Q5 DNA polymerase, HiScribe T7 High Yield RNA kit, and restriction endonucleases were purchased from New England Biolabs (Ipswich, MA, USA). E.Z.N.A. Plasmid Mini and Midi Kits were purchased from Omega Bio-Tek (Norcross, GA, USA), and QIAquick PCR Purification Kits were purchased from QIAGEN (Valencia, CA, USA).

### Molecular Cloning

*Escherichia coli* K-12 DH10B (New England Biolabs, Ipswich, MA, USA) was used for plasmid cloning. The plasmid pJL1, with a ColE1 origin and kanamycin resistance cassette, was used as the backbone vector for all plasmids.^43^

All constructs were assembled with inverse PCR or Gibson assembly^22^. LB medium composed of 10 g/L NaCl, 5 g/L yeast extract, and 10 g/L tryptone was used for all cell growth during cloning steps. Kanamycin (33 µg/mL) was used for appropriate antibiotic selection. The coding sequences for the *eutR*, *metR*, *ulaR*, *ulaA, ulaB*, *ulaC, ZntR, and Zur* genes and related promoters were isolated from *E. coli* K-12 DH10B genomic DNA. ^14^ All plasmid sequences are described in the Supplementary Information document.

### Preparation of cellular lysate for cell-free reactions

Unless otherwise specified, cellular lysate for all experiments was prepared as previously described^23^. *E. coli* BL21 Star (DE3) *ΔlacZ* cells were grown in 2x YTP medium at 37°C and 180 rpm to an OD_600_ of 1.7, which corresponded with the mid-exponential growth phase. Expression of T7 RNA polymerase was induced with 0.4 mM isopropyl β-d-1-thiogalactopyranoside (IPTG) around OD_600_ 0.6. Cells were then centrifuged at 2700 ×g and washed three times with S30A buffer (50 mM tris, 14 mM magnesium glutamate, 60 mM potassium glutamate, and 2 mM dithiothreitol, pH 7.7.) After the final centrifugation, the wet cell mass was determined, and cells were resuspended in 1 mL of S30A buffer per 1 g of wet cell mass. The cellular resuspension was divided into 1 mL aliquots. Cells were lysed using a Q125 Sonicator (Qsonica, Newton, CT, USA) at a frequency of 20 kHz, and at 50% of amplitude. Cells were sonicated on ice with three cycles of 10 seconds on, 10 seconds off, delivering approximately 300 J, at which point the cells appeared visibly lysed. An additional 4 mM of dithiothreitol was added to each tube, and the sonicated mixture was then centrifuged at 12,000 ×g and 4°C for 10 minutes. The supernatant was removed, divided into 1 mL aliquots, and incubated at 37°C and 220 rpm for 80 minutes. After this runoff reaction, the cellular lysate was centrifuged at 12,000 ×g and 4°C for 10 minutes. The supernatant was removed and loaded into a 10 kDa MWCO dialysis cassette (Thermo Fisher). Lysate was dialyzed in 1 L of S30B buffer (14 mM magnesium glutamate, 60 mM potassium glutamate, 1 mM dithiothreitol, pH 8.2) at 4°C for 3 hours. Dialyzed lysate was removed and centrifuged at 12,000 ×g and 4°C for 10 minutes. The supernatant was removed, aliquoted, and stored at −80°C for future use.

Cellular lysate for the homocysteine and vitamin B_12_ biosensors was prepared as described above but was not induced with IPTG during cell growth. The lysate enriched with phosphotransferase proteins (UlaABC) were prepared from *E. coli* BL21 Star (DE3) *ΔlacZ* transformants carrying a plasmid expressing UlaABC from a P_BAD, weak_ promoter. Expression of UlaABC was induced by adding arabinose at a concentration of 1 mM around OD_600_ 0.6. This lysate was also induced to increase expression of T7 RNA polymerase.

### Preparation of non-coding RNA

RNA for sfGFP without an RBS was transcribed from linear DNA templates using HiScribe T7 High Yield RNA synthesis kit according to the manufacturer’s protocol (New England Biolabs). Following RNA synthesis, Dnase I (Zymo Research) was added to degrade the linear DNA template. The RNA products were then purified using an RNA Clean and Concentrator kit (Zymo Research) according to the manufacturer’s protocol. Following purification, RNA concentration was measured on a Nanodrop 2000, sub-aliquoted to reduce freeze-thaw cycles, and stored −80 °C.

### Cell-free reactions

Cell-free reactions for all experiments were run as previously described^24^. Each cell-free reaction contained 0.85 mM each of GTP, UTP, and CTP, in addition to 1.2 mM ATP, 34 μg/mL of folinic acid, 170 μg/mL *E. coli* tRNA mixture, 130 mM potassium glutamate, 10 mM ammonium glutamate, 12 mM magnesium glutamate, 2 mM each of the 20 standard amino acids, 0.33 mM nicotine adenine dinucleotide (NAD), 0.27 mM coenzyme-A (CoA), 1.5 mM spermidine, 1 mM putrescine, 4 mM sodium oxalate, 33 mM phosphoenolpyruvate (PEP), 27 % (v/v) cell extract, and plasmid concentrations specified for each experiment. In reactions producing LacZ (β-galactosidase), CPRG was added to a final concentration of 0.6 mg/mL.

Reactions were run in 10 μL volumes in 384-well small volume plates (Greiner Bio-One), and a clear adhesive film was used to cover the plate and prevent evaporation. Unless otherwise noted, Figures 1-3 plates were incubated for 4h at 37 °C and Figure 5 was incubated for 3h at 37 °C. Fluorescence was measured with a plate reader (Synergy4, BioTek). Time-course experiments were run directly in the plate reader. Excitation and emission wavelengths for sfGFP were 485 and 510 nm, respectively.

### Data Processing and Statistical Analysis

Fluorescence values are background subtracted using the fluorescence of a reaction containing no plasmid. The limit of detection was determined by comparing the fluorescence values of reactions with a specified analyte concentration with those of reactions with no added analyte using a two-tailed student’s t-test assuming equal variance. The lowest concentration of analyte that yielded a p-value below 0.05 for that concentration and all higher concentrations was considered the limit of detection.

The dynamic range was computed as the fold change in fluorescence between the reaction containing no analyte and the reaction that generates the largest fluorescence value.

To one-dimensonalize crosstalk’s effects on LacZ expression, the area under the curve (AUC) was calculated for both leak and activation, with and without the non-regulatory plasmid. CMT effects were quantified as ΔAUC=AUC_NonRegulatory_-AUC_Base_. To quantify the total effect of crosstalk on apparent biosensor performance, the area between the activation and leak curves (ABC) under both conditions was calculated, with ΔABC=ABC_NonRegulatory_-ABC_BaseCase_. For both ΔAUC and ΔABC a value above 1 corresponds with positive crosstalk and a value below 1 corresponds with negative crosstalk (Figure S4-S5).

## Supporting Information

Supplementary Figures S1: Raw data for Figure 1A. S2: Raw data for Figure 3A and 3B. S3: Crosstalk in the vitamin C biosensor generated by expression of transcription factors from plasmids prepped on different days. S4-S5: Visual representation of data analysis used in Figure 4. S6: Raw data from Figure 4C. S7: Inefficient inversion of colorimetric reporter at high plasmid concentrations.

Supplementary Table S1: Annotated sequences of all plasmids.

## Author Contributions

Conceptualization: AP, FP, and MPS; Investigation: AP, FP, SB, and DG; Formal Analysis: AP, FP, SB, and DG; Writing – Original Draft: AP and FP; Writing – Review & Editing: AP, FP, and MPS; Visualization: AP and FP; Supervision: MPS; Funding Acquisition: MPS.

## Acknowledgments

**Funding:** This work was supported by the National Science Foundation (EF-2319391 and DGE-2039665) and the National Institutes of Health (R01EB034301, R35GM149268, and F99DK143562).

## Conflicts of Interest

None

## Notes

### Competing Interest Statement

The authors have declared no competing interest.

