## Supplementary Information for "Harnessing Plasmid Crosstalk in Transcription Factor Mediated Cell-Free Biosensors"

**Harnessing Plasmid Crosstalk as a Design Tool in *Escherichia coli*-Based Cell-Free Biosensors**

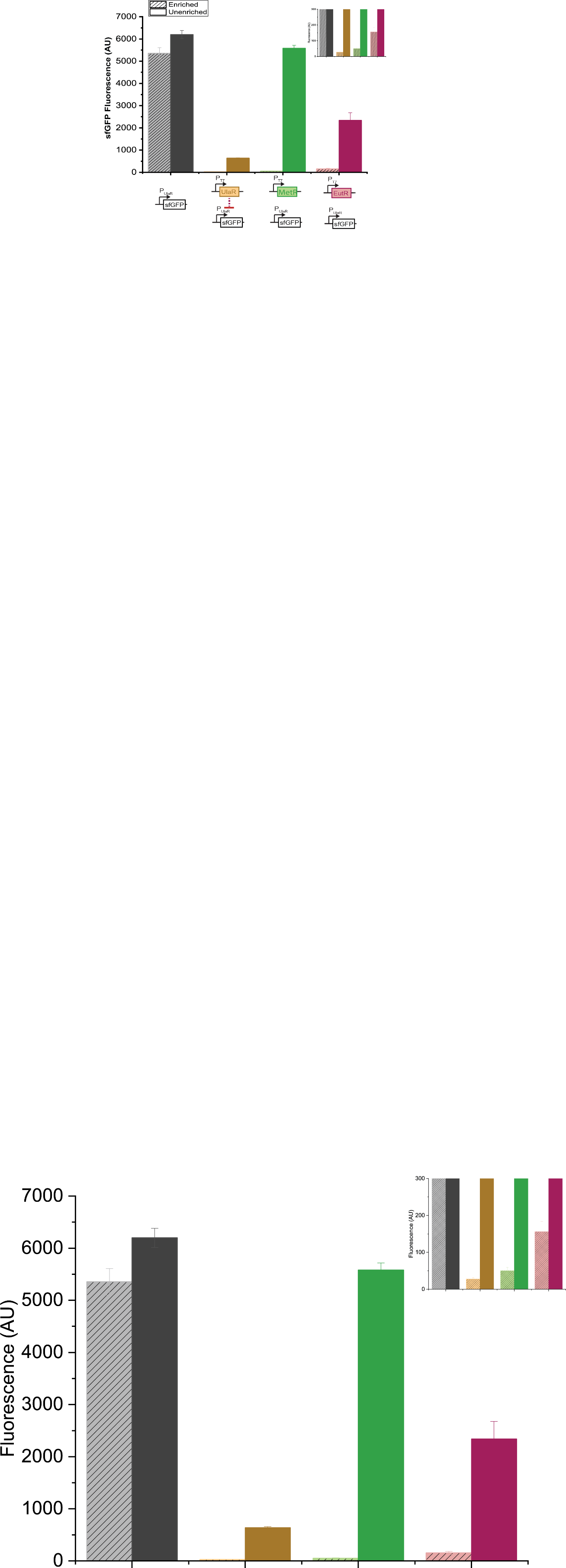

**Figure S1.** Crosstalk in the vitamin C biosensor generated by expression of transcription factors. This figure depicts the raw data shown in Figure 1B. Data were collected after 4 hours of incubation at 37°C. Background fluorescence was subtracted from all samples and error bars indicate the standard deviation of three technical replicates.

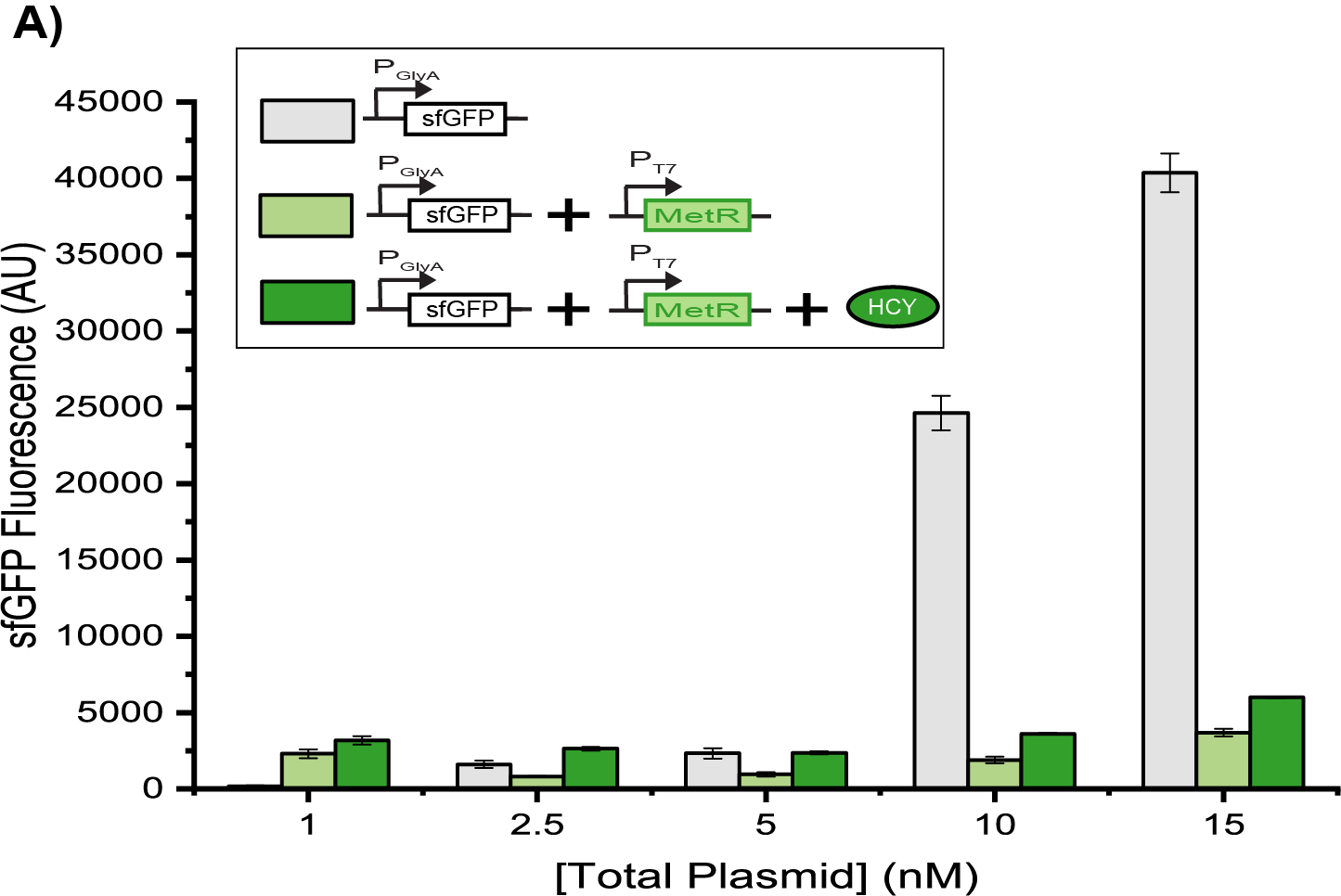

**Figure S2.** Crosstalk in the homocysteine biosensor generated by expression of MetR. This figure depicts the raw data shown Figure 3. Data were collected after 4 hours of incubation at 37°C. Background fluorescence was subtracted from all samples and error bars indicate the standard deviation of three technical replicates.

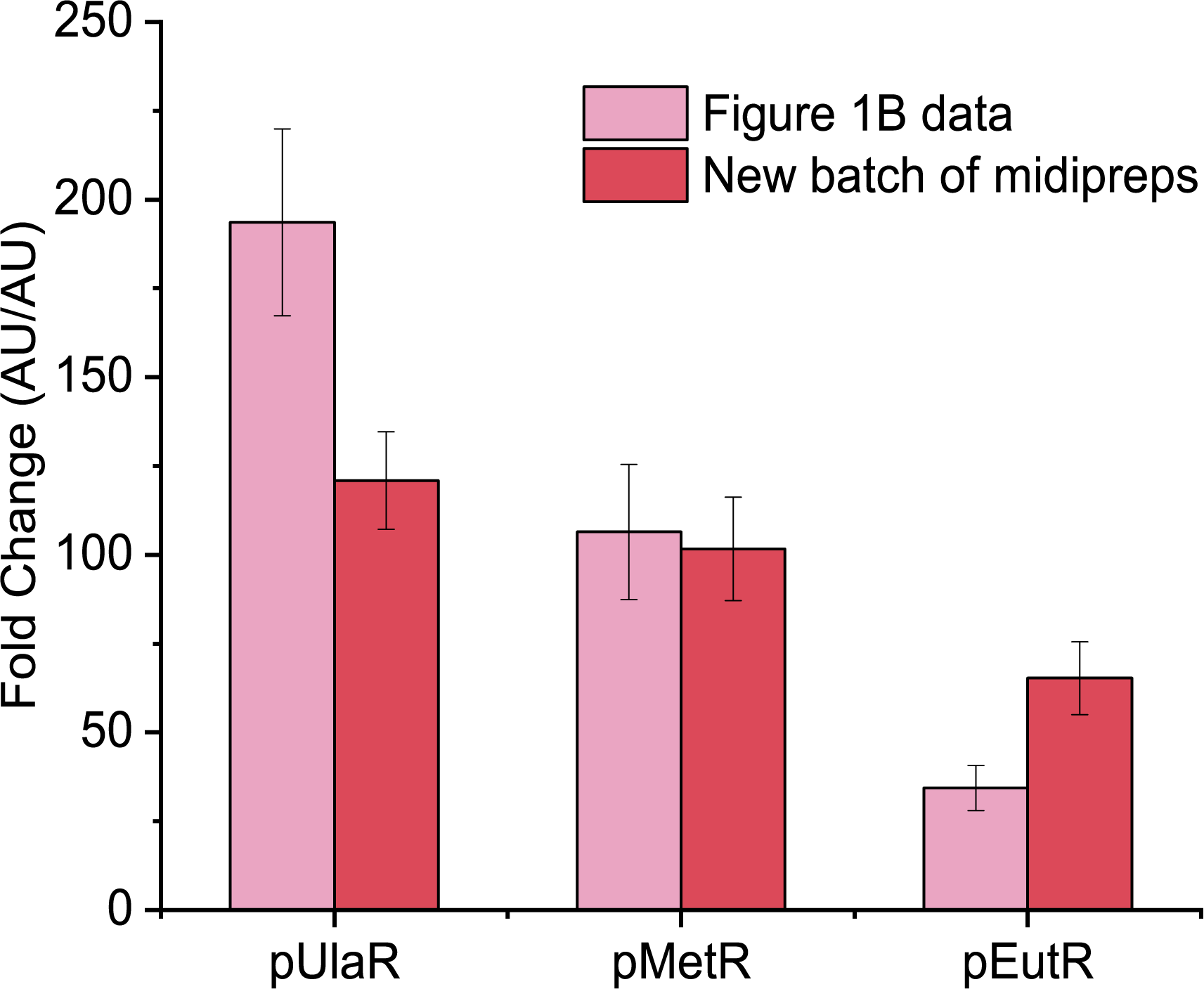

**Figure S3.** Crosstalk in the vitamin C biosensor generated by expression of transcription factors. This figure reproduces the experimental design of Figure 1B, but with freshly prepared plasmids. Data were collected after 4 hours of incubation at 37°C. Background fluorescence was subtracted from all samples and error bars indicate the standard deviation of three technical replicates.

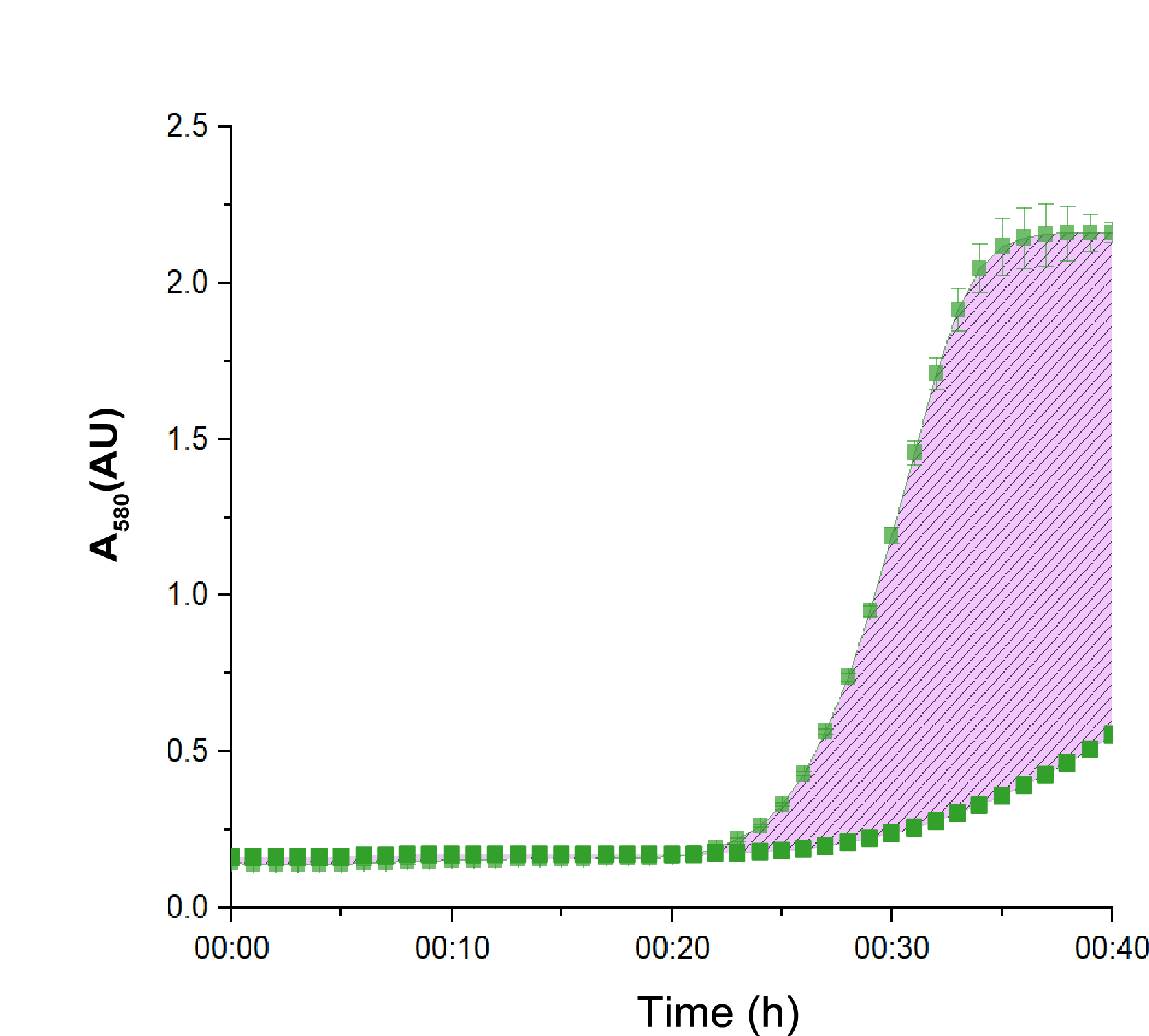

**Figure S4.** Visual depiction of area-between-curve calculations. Top line is the activation condition (with zinc). Bottom line is the non-induced condition (without zinc). The purple crosshatched sections visually depict the ABC. When calculating ΔABC, the ABCs were calculated for each individual condition (base vs crosstalk) and then subtracted. Data were collected during incubation at 37°C. Error bars indicate the standard deviation of three technical replicates.

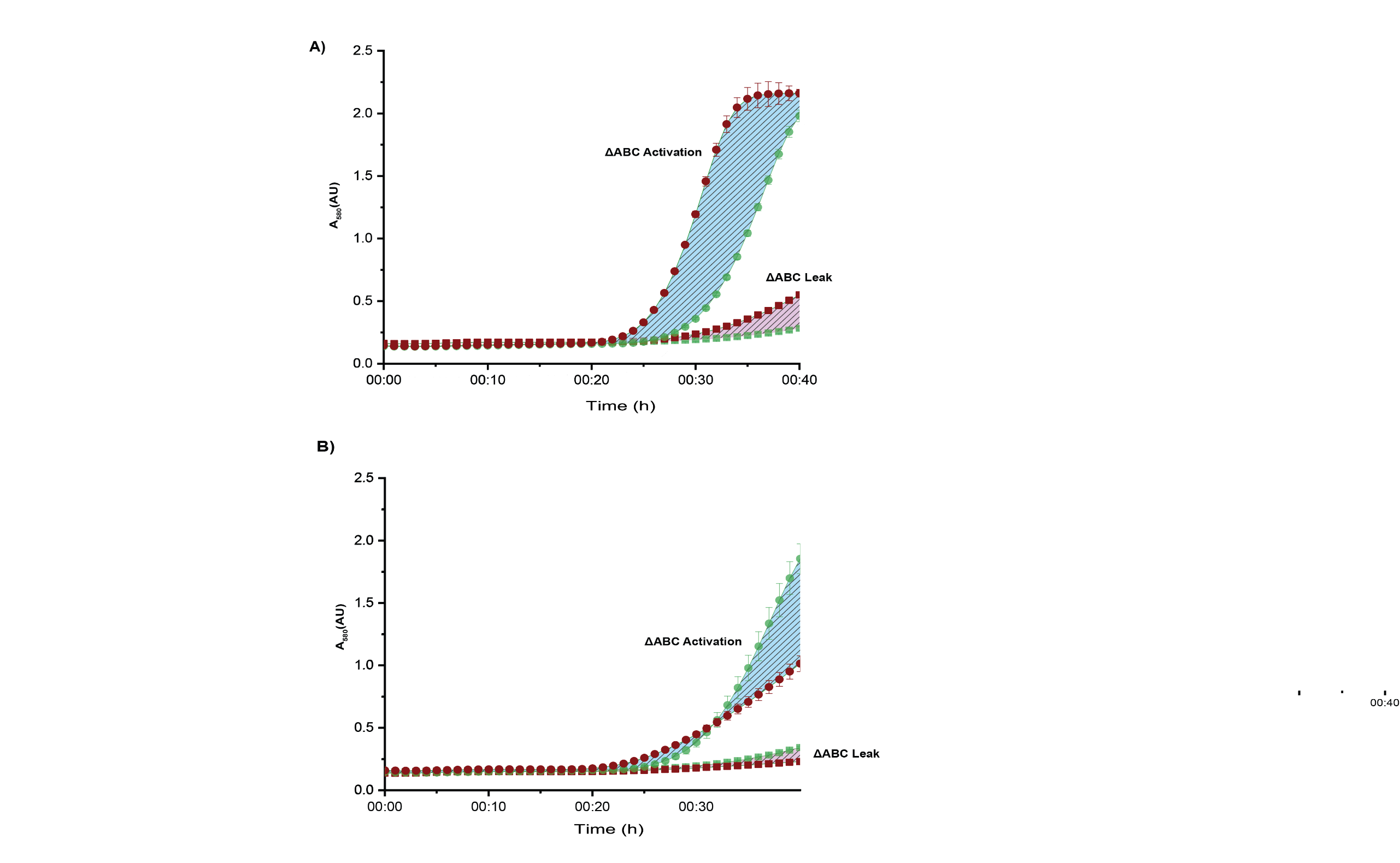

**Figure S5.** Visual depiction of the data analysis as shown in Figure 4C. In all panels, green lines represent the base case and red lines depict the expression when a non-regulatory plasmid is added. Dotted lines are the activation condition (with zinc). Square lines are the non-induced condition (without zinc). The blue crosshatched sections visually depict ΔAUC for activation and the purple crosshatched sections visually depict the ΔAUC for leak. **(A)** Example of positive crosstalk, resulting in a positive ΔAUC for activation and leak. **(B)** Example of negative crosstalk, resulting in a negative ΔAUC for activation and leak. Data were collected during incubation at 37°C. Error bars indicate the standard deviation of three technical replicates.

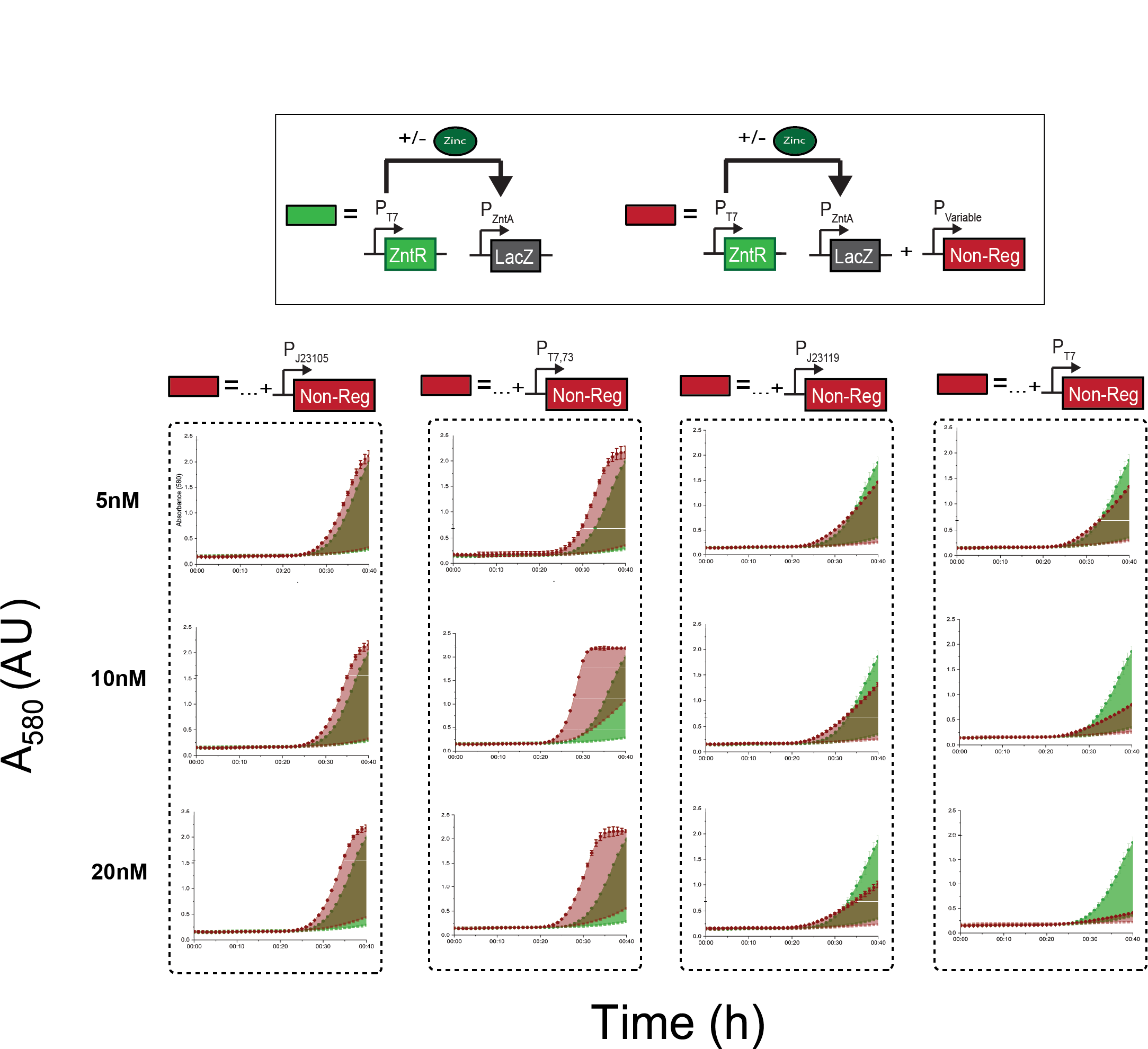

**Figure S6.** Crosstalk in the zinc activator biosensor generated by expression of a non-regulatory plasmid. This figure depicts the raw data shown in Figure 4C-E. Data were collected during incubation at 37°C. Error bars indicate the standard deviation of three technical replicates.

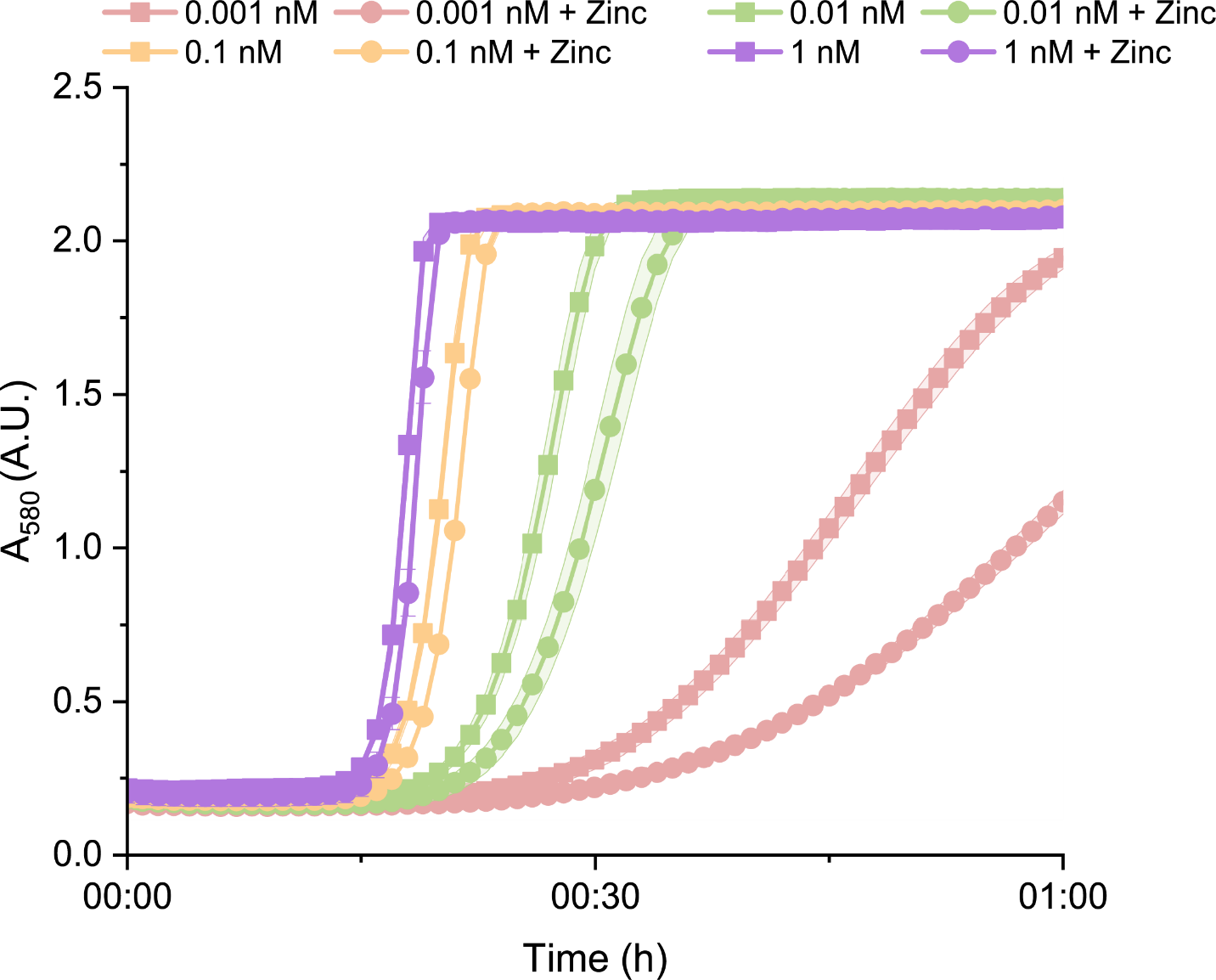

**Figure S7.** Comparison of inversion efficiency at different concentrations of pT7LacZ. At high concentrations of plasmid, inversion does not occur. Data were collected during incubation at 37°C. Error bars indicate the standard deviation of three technical replicates.

**Table 1**. Annotated sequences of all plasmids used in this paper.

| pUlaAsfGFP | Plasmid encoding sfGFP expression under P_UlaA_ |
| --- | --- |
| P_UlaG_-P_UlaA_ Stability hairpin RBS sfGFP Terminator Kanamycin resistance cassette ColE1 origin  agatcaaaggatcttcttgagatcctttttttctgcgcgtaatctgctgcttgcaaacaaaaaaaccaccgctaccagcggtggtttgtttgccggatcaagagctaccaactctttttccgaaggtaactggcttcagcagagcgcagataccaaatactgttcttctagtgtagccgtagttaggccaccacttcaagaactctgtagcaccgcctacatacctcgctctgctaatcctgttaccagtggctgctgccagtggcgataagtcgtgtcttaccgggttggactcaagacgatagttaccggataaggcgcagcggtcgggctgaacggggggttcgtgcacacagcccagcttggagcgaacgacctacaccgaactgagatacctacagcgtgagctatgagaaagcgccacgcttcccgaagggagaaaggcggacaggtatccggtaagcggcagggtcggaacaggagagcgcacgagggagcttccagggggaaacgcctggtatctttatagtcctgtcgggtttcgccacctctgacttgagcgtcgatttttgtgatgctcgtcaggggggcggagcctatggaaaaacgccagcaacgcgatcccgcgaaatCGCCATTTACCTTCATGATAGTTCAATTCGAATCAATATGTGATTGGTTTTGATTAATCCTGACACTATTTTTTCAGGAAGGCAATGACCATTTTTTGACTTTTTGCCAGGGAAGTTGTTGTTGATTTTGAGTATGGAAAGATTTAATGGAATGTGTAATTCAATTAACTAAATGAATTTAAATGGATAATTGTTTCGTTGTGTGAATCCCACTCTATCCATGTGGAATTATTTGCGGGTCGCGTCACATTTAATCATAAATAATCTTGTTGTGATTACTTTTGAAAATTAGAGTGAGTGCACAACATTCCGGGTGTGTGGAATACCCGGTTACCTCTTCTTCAGGAGATCGTTccacaacggtttccctctagaaataattttgtttaactttaagaaggagatatacatATGAGCAAAGGTGAAGAACTGTTTACCGGCGTTGTGCCGATTCTGGTGGAACTGGATGGCGATGTGAACGGTCACAAATTCAGCGTGCGTGGTGAAGGTGAAGGCGATGCCACGATTGGCAAACTGACGCTGAAATTTATCTGCACCACCGGCAAACTGCCGGTGCCGTGGCCGACGCTGGTGACCACCCTGACCTATGGCGTTCAGTGTTTTAGTCGCTATCCGGATCACATGAAACGTCACGATTTCTTTAAATCTGCAATGCCGGAAGGCTATGTGCAGGAACGTACGATTAGCTTTAAAGATGATGGCAAATATAAAACGCGCGCCGTTGTGAAATTTGAAGGCGATACCCTGGTGAACCGCATTGAACTGAAAGGCACGGATTTTAAAGAAGATGGCAATATCCTGGGCCATAAACTGGAATACAACTTTAATAGCCATAATGTTTATATTACGGCGGATAAACAGAAAAATGGCATCAAAGCGAATTTTACCGTTCGCCATAACGTTGAAGATGGCAGTGTGCAGCTGGCAGATCATTATCAGCAGAATACCCCGATTGGTGATGGTCCGGTGCTGCTGCCGGATAATCATTATCTGAGCACGCAGACCGTTCTGTCTAAAGATCCGAACGAAAAAGGCACGCGGGACCACATGGTTCTGCACGAATATGTGAATGCGGCAGGTATTACGTGGAGCCATCCGCAGTTCGAAAAATAAgtcgaccggctgctaacaaagcccgaaaggaagctgagttggctgctgccaccgctgagcaataactagcataaccccttggggcctctaaacgggtcttgaggggttttttgctgaaagccaattctgattagaaaaactcatcgagcatcaaatgaaactgcaatttattcatatcaggattatcaataccatatttttgaaaaagccgtttctgtaatgaaggagaaaactcaccgaggcagttccataggatggcaagatcctggtatcggtctgcgattccgactcgtccaacatcaatacaacctattaatttcccctcgtcaaaaataaggttatcaagtgagaaatcaccatgagtgacgactgaatccggtgagaatggcaaaagcttatgcatttctttccagacttgttcaacaggccagccattacgctcgtcatcaaaatcactcgcatcaaccaaaccgttattcattcgtgattgcgcctgagcgagacgaaatacgcgatcgctgttaaaaggacaattacaaacaggaatcgaatgcaaccggcgcaggaacactgccagcgcatcaacaatattttcacctgaatcaggatattcttctaatacctggaatgctgttttcccggggatcgcagtggtgagtaaccatgcatcatcaggagtacggataaaatgcttgatggtcggaagaggcataaattccgtcagccagtttagtctgaccatctcatctgtaacatcattggcaacgctacctttgccatgtttcagaaacaactctggcgcatcgggcttcccatacaatcgatagattgtcgcacctgattgcccgacattatcgcgagcccatttatacccatataaatcagcatccatgttggaatttaatcgcggcttcgagcaagacgtttcccgttgaatatggctcataacaccccttgtattactgtttatgtaagcagacagttttattgttcatgatgatatatttttatcttgtgcaatgtaacatcagagattttgagacacaacgtg | |

| pUlaR | Plasmid encoding UlaR expression under P_T7_ |
| --- | --- |
| P_T7_ Stability hairpin RBS *ulaR* Terminator Kanamycin resistance cassette ColE1 origin  agatcaaaggatcttcttgagatcctttttttctgcgcgtaatctgctgcttgcaaacaaaaaaaccaccgctaccagcggtggtttgtttgccggatcaagagctaccaactctttttccgaaggtaactggcttcagcagagcgcagataccaaatactgttcttctagtgtagccgtagttaggccaccacttcaagaactctgtagcaccgcctacatacctcgctctgctaatcctgttaccagtggctgctgccagtggcgataagtcgtgtcttaccgggttggactcaagacgatagttaccggataaggcgcagcggtcgggctgaacggggggttcgtgcacacagcccagcttggagcgaacgacctacaccgaactgagatacctacagcgtgagctatgagaaagcgccacgcttcccgaagggagaaaggcggacaggtatccggtaagcggcagggtcggaacaggagagcgcacgagggagcttccagggggaaacgcctggtatctttatagtcctgtcgggtttcgccacctctgacttgagcgtcgatttttgtgatgctcgtcaggggggcggagcctatggaaaaacgccagcaacgcgatcccgcgaaattaatacgactcactatagggagaccacaacggtttccctctagaaataattttgtttaactttaagaaggagatatacatatgACTGAAGCACAAAGACATCAAATCCTCCTGGAAATGCTCGCACAATTGGGCTTTGTGACCGTTGAGAAAGTCGTTGAGCGTCTGGGAATTTCGCCTGCCACTGCGCGACGCGATATCAATAAACTTGACGAAAGCGGCAAACTGAAAAAAGTGCGCAATGGCGCAGAAGCTATTACCCAACAGCGCCCGCGCTGGACGCCGATGAATCTGCATCAGGCGCAGAATCACGATGAAAAAGTACGTATCGCTAAAGCGGCCTCGCAGCTGGTTAATCCGGGCGAAAGCGTAGTCATCAACTGCGGCTCCACCGCGTTTCTGCTTGGGCGGGAAATGTGTGGCAAGCCAGTGCAAATCATCACTAATTATCTACCGCTGGCAAATTACCTGATCGATCAAGAACATGACAGCGTGATCATTATGGGCGGACAGTACAACAAAAGTCAGTCCATCACTTTAAGCCCGCAGGGCAGCGAAAACAGTCTCTATGCCGGGCACTGGATGTTTACCAGCGGAAAAGGGCTGACCGCAGAAGGGTTGTATAAAACCGATATGCTGACAGCAATGGCAGAGCAGAAGATGCTGAGCGTGGTAGGGAAACTGGTGGTACTGGTTGATAGCAGTAAGATTGGCGAACGCGCGGGAATGCTTTTTAGCCGTGCCGATCAAATCGATATGCTTATCACCGGCAAAAATGCTAACCCGGAAATCCTGCAACAACTGGAAGCGCAAGGGGTCAGCATTCTGCGTGTTtaagtcgaccggctgctaacaaagcccgaaaggaagctgagttggctgctgccaccgctgagcaataactagcataaccccttggggcctctaaacgggtcttgaggggttttttgctgaaagccaattctgattagaaaaactcatcgagcatcaaatgaaactgcaatttattcatatcaggattatcaataccatatttttgaaaaagccgtttctgtaatgaaggagaaaactcaccgaggcagttccataggatggcaagatcctggtatcggtctgcgattccgactcgtccaacatcaatacaacctattaatttcccctcgtcaaaaataaggttatcaagtgagaaatcaccatgagtgacgactgaatccggtgagaatggcaaaagcttatgcatttctttccagacttgttcaacaggccagccattacgctcgtcatcaaaatcactcgcatcaaccaaaccgttattcattcgtgattgcgcctgagcgagacgaaatacgcgatcgctgttaaaaggacaattacaaacaggaatcgaatgcaaccggcgcaggaacactgccagcgcatcaacaatattttcacctgaatcaggatattcttctaatacctggaatgctgttttcccggggatcgcagtggtgagtaaccatgcatcatcaggagtacggataaaatgcttgatggtcggaagaggcataaattccgtcagccagtttagtctgaccatctcatctgtaacatcattggcaacgctacctttgccatgtttcagaaacaactctggcgcatcgggcttcccatacaatcgatagattgtcgcacctgattgcccgacattatcgcgagcccatttatacccatataaatcagcatccatgttggaatttaatcgcggcttcgagcaagacgtttcccgttgaatatggctcataacaccccttgtattactgtttatgtaagcagacagttttattgttcatgatgatatatttttatcttgtgcaatgtaacatcagagattttgagacacaacgtg | |

| pMetR | Plasmid encoding MetR expression under P_T7_ |
| --- | --- |
| P_T7_ Stability hairpin RBS *metR* Terminator Kanamycin resistance cassette ColE1 origin  agatcaaaggatcttcttgagatcctttttttctgcgcgtaatctgctgcttgcaaacaaaaaaaccaccgctaccagcggtggtttgtttgccggatcaagagctaccaactctttttccgaaggtaactggcttcagcagagcgcagataccaaatactgttcttctagtgtagccgtagttaggccaccacttcaagaactctgtagcaccgcctacatacctcgctctgctaatcctgttaccagtggctgctgccagtggcgataagtcgtgtcttaccgggttggactcaagacgatagttaccggataaggcgcagcggtcgggctgaacggggggttcgtgcacacagcccagcttggagcgaacgacctacaccgaactgagatacctacagcgtgagctatgagaaagcgccacgcttcccgaagggagaaaggcggacaggtatccggtaagcggcagggtcggaacaggagagcgcacgagggagcttccagggggaaacgcctggtatctttatagtcctgtcgggtttcgccacctctgacttgagcgtcgatttttgtgatgctcgtcaggggggcggagcctatggaaaaacgccagcaacgcgatcccgcgaaattaatacgactcactatagggagaccacaacggtttccctctagaaataattttgtttaactttaagaaggagatatacatatgatcgaagtaaaacacctgaaaacgctacaagcgttgcggaactgcggctcgctcgcagccgctgcggcgacgttgcatcagacgcaatccgccctgtctcaccagtttagcgatctggaacaacgccttggcttccggctatttgtgcgtaagagccagccgctacgctttacaccgcagggagaaatcctgttgcaactggcaaaccaggtactgccgcaaattagccaggccctgcaagcctgcaatgaaccgcagcagacgcgtctgcgcattgccattgagtgccatagctgtattcagtggctgacacccgcgttagaaaatttccataagaactggccgcaggtagagatggattttaaatcgggcgtgacatttgacccgcagcccgccttgcaacagggagagctggatctggtaatgacgtccgatattctgccgcgcagtggcctgcattattcgccgatgttcgactatgaagtgcgtctggtgttagcacctgaccatccactggcggcgaaaacgcgaattacaccggaagatctcgccagcgagacgctattaatttatccggtgcagcgtagtcgactggatgtctggcggcattttcttcagccggcaggcgtcagcccgtcactgaaaagcgtcgataacaccttattgttgattcagatggttgccgcgcggatgggtattgccgcgctaccgcattgggtagtagagagttttgagcgccagggtctggtggtgaccaaaaccctgggcgaaggcttgtggagccgactgtacgccgccgtgcgcgatggcgagcagcgtcagccagttacggaagcgtttattcgctcagcgcgcaatcacgcctgcgatcatctgccgtttgtgaagagcgcggagcgacccacttacgatgcacccacagtgaggccaggatcaccagcgcgcctgtaagtcgaccggctgctaacaaagcccgaaaggaagctgagttggctgctgccaccgctgagcaataactagcataaccccttggggcctctaaacgggtcttgaggggttttttgctgaaagccaattctgattagaaaaactcatcgagcatcaaatgaaactgcaatttattcatatcaggattatcaataccatatttttgaaaaagccgtttctgtaatgaaggagaaaactcaccgaggcagttccataggatggcaagatcctggtatcggtctgcgattccgactcgtccaacatcaatacaacctattaatttcccctcgtcaaaaataaggttatcaagtgagaaatcaccatgagtgacgactgaatccggtgagaatggcaaaagcttatgcatttctttccagacttgttcaacaggccagccattacgctcgtcatcaaaatcactcgcatcaaccaaaccgttattcattcgtgattgcgcctgagcgagacgaaatacgcgatcgctgttaaaaggacaattacaaacaggaatcgaatgcaaccggcgcaggaacactgccagcgcatcaacaatattttcacctgaatcaggatattcttctaatacctggaatgctgttttcccggggatcgcagtggtgagtaaccatgcatcatcaggagtacggataaaatgcttgatggtcggaagaggcataaattccgtcagccagtttagtctgaccatctcatctgtaacatcattggcaacgctacctttgccatgtttcagaaacaactctggcgcatcgggcttcccatacaatcgatagattgtcgcacctgattgcccgacattatcgcgagcccatttatacccatataaatcagcatccatgttggaatttaatcgcggcttcgagcaagacgtttcccgttgaatatggctcataacaccccttgtattactgtttatgtaagcagacagttttattgttcatgatgatatatttttatcttgtgcaatgtaacatcagagattttgagacacaacgtg | |

| pEutR | Plasmid encoding EutR expression under P_T7_ |
| --- | --- |
| P_T7_ Stability hairpin RBS *eutR* Terminator Kanamycin resistance cassette ColE1 origin  agatcaaaggatcttcttgagatcctttttttctgcgcgtaatctgctgcttgcaaacaaaaaaaccaccgctaccagcggtggtttgtttgccggatcaagagctaccaactctttttccgaaggtaactggcttcagcagagcgcagataccaaatactgttcttctagtgtagccgtagttaggccaccacttcaagaactctgtagcaccgcctacatacctcgctctgctaatcctgttaccagtggctgctgccagtggcgataagtcgtgtcttaccgggttggactcaagacgatagttaccggataaggcgcagcggtcgggctgaacggggggttcgtgcacacagcccagcttggagcgaacgacctacaccgaactgagatacctacagcgtgagctatgagaaagcgccacgcttcccgaagggagaaaggcggacaggtatccggtaagcggcagggtcggaacaggagagcgcacgagggagcttccagggggaaacgcctggtatctttatagtcctgtcgggtttcgccacctctgacttgagcgtcgatttttgtgatgctcgtcaggggggcggagcctatggaaaaacgccagcaacgcgatcccgcgaaattaatacgactcactatagggagaccacaacggtttccctctagaaataattttgtttaactttaagaaggagatatacat atgaaaaagacccgtacagccaatttgcaccatctttatcatgaacccttacccgaaaacctgaagctcacgccgaaggtcgaagtggataatgttcatcaacgacagacaacggatgtctatgaacatgctttaacgattaccgcctggcagcagatttacgatcagctgcatccgggcaagtttcatggtgaatttacggaaattctactcgatgatattcaggtttttcgtgaatacaccggtctggcgctgcgtcagtcgtgcctggtctggccgaactcgttctggtttggcattccggcgacgcgcggtgagcagggatttatcggttcgcaatgtctgggaagcgcggaaatcgccacccgccctggtggcactgaatttgaactgagcacgccggatgattacacgatcctgggcgtggtgctttctgaagatgtcatcacccggcaggctaactttttgcataacccggatcgggtattacatatgttgcgtaaccagtcggcgctggaagtgaaagagcagcataaagccgcgctgtggggctttgtccaacaggcgctggcgacgttttgcgagaatccggaaaatctccatcagccagcagtgcgaaaagtgctgggggataatttgctaatggcgatgggggccatgctggaagaagcgcaaccaatggtgacggcggaaagcatcagtcatcagagttaccgtcgattgctttcccgcgcccgtgaatatgtgctggaaaacatgtccgaaccggtgacggtgctggatttgtgtaatcaactgcatgtcagccgccgcacgctacaaaacgcgtttcacgctattttaggcattggcccgaacgcgtggctgaaacgcattcgcctgaacgccgtacgccgcgaactgataagtccgtggtcgcaaagtatgacggtaaaagacgccgccatgcagtggggattctggcatctggggcaatttgccacggattaccagcagctgttttccgagaagccgtcactgacgctgcatcagcggatgcgggagtgggggtgagtcgaccggctgctaacaaagcccgaaaggaagctgagttggctgctgccaccgctgagcaataactagcataaccccttggggcctctaaacgggtcttgaggggttttttgctgaaagccaattctgattagaaaaactcatcgagcatcaaatgaaactgcaatttattcatatcaggattatcaataccatatttttgaaaaagccgtttctgtaatgaaggagaaaactcaccgaggcagttccataggatggcaagatcctggtatcggtctgcgattccgactcgtccaacatcaatacaacctattaatttcccctcgtcaaaaataaggttatcaagtgagaaatcaccatgagtgacgactgaatccggtgagaatggcaaaagcttatgcatttctttccagacttgttcaacaggccagccattacgctcgtcatcaaaatcactcgcatcaaccaaaccgttattcattcgtgattgcgcctgagcgagacgaaatacgcgatcgctgttaaaaggacaattacaaacaggaatcgaatgcaaccggcgcaggaacactgccagcgcatcaacaatattttcacctgaatcaggatattcttctaatacctggaatgctgttttcccggggatcgcagtggtgagtaaccatgcatcatcaggagtacggataaaatgcttgatggtcggaagaggcataaattccgtcagccagtttagtctgaccatctcatctgtaacatcattggcaacgctacctttgccatgtttcagaaacaactctggcgcatcgggcttcccatacaatcgatagattgtcgcacctgattgcccgacattatcgcgagcccatttatacccatataaatcagcatccatgttggaatttaatcgcggcttcgagcaagacgtttcccgttgaatatggctcataacaccccttgtattactgtttatgtaagcagacagttttattgttcatgatgatatatttttatcttgtgcaatgtaacatcagagattttgagacacaacgtg | |

| pGlyAsfGFP | Plasmid encoding sfGFP expression under P_GlyA_ |
| --- | --- |
| P_GlyA_ Stability hairpin RBS sfGFP Terminator Kanamycin resistance cassette ColE1 origin  agatcaaaggatcttcttgagatcctttttttctgcgcgtaatctgctgcttgcaaacaaaaaaaccaccgctaccagcggtggtttgtttgccggatcaagagctaccaactctttttccgaaggtaactggcttcagcagagcgcagataccaaatactgttcttctagtgtagccgtagttaggccaccacttcaagaactctgtagcaccgcctacatacctcgctctgctaatcctgttaccagtggctgctgccagtggcgataagtcgtgtcttaccgggttggactcaagacgatagttaccggataaggcgcagcggtcgggctgaacggggggttcgtgcacacagcccagcttggagcgaacgacctacaccgaactgagatacctacagcgtgagctatgagaaagcgccacgcttcccgaagggagaaaggcggacaggtatccggtaagcggcagggtcggaacaggagagcgcacgagggagcttccagggggaaacgcctggtatctttatagtcctgtcgggtttcgccacctctgacttgagcgtcgatttttgtgatgctcgtcaggggggcggagcctatggaaaaacgccagcaacgcgatcccgcgaaat GCCGCATGGAACCAGTTCACTTGTGCTGTGTGGCCTGCTTTTGCCAGCGTGTCGAGCATTGCCAGCATTGGCGTTTGACCAACACCGGCAGAGATTAACGTCACTGGTGTGTCATCTGCGACAGCCATAAAGAAATCACCTGCCGGAGCGACCAGTTTCACGACATCGCCAACATTGGCGTGATTGTGCAACCAGTTGGATACCTGCCCACCCTCTTCGCGTTTCACCGCAATACGATAGCCTTTGCCATCCGGTTTGCGAGTCAAAGAGTACTGACGAATTTCCTGATGTGGGAAACCTTCCGGCTTCAGCCAGACGCCGAGATATTGCCCCGGACGGTATTCTGCCACTGCGCCACCGTCGACCGGCTCCAGTTCGAAGCTGGTGATAAGCGCGCTGCGCGGTGTTTTAGCCACAATGCGGAAATCGCGAGTACCTTCCCAACCACCGGCTTTGCTGGCGTTTTCGTTATAGATTTCCGCCTCGCGATTGATAAATACATTAGCCAGTACACCATAGGCTTTACCCCACGCGTCCAGCACTTCCTGCCCCGGGCTGAACATTTCGTCCAGCGTTGCCAACAGGTGTTCACCGACGATGTTGTACTGTTCCGGTTTGATCTGGAAGCTGGTGTGCTTCTGCGCGATTTTTTCTACCGCTGGCAGCAGCGCAGGCAGGTTTTCAATATTACTGGCGTAGGCGGCAATAGCGTTAAACAGGGCTTCACGTTGATCGCCATTACGCTGGTTACTCATGTTAAAAATTTCTTTGAGTTCTGGGTTATGAGTAAACATACGGTCGTAGAAATGGGCGGTTAACTTTGGCCCCGTTTCCACCAGTAAAGGGATGGTGGCTTTTACTGTAGCGATGGTTTGAGCGTCAAGCATATGGTCTTCCTTTTTTTGCATCTTAATTGATGTATCTCAAATGCATCTTATAAAAAATAGCCCTGCAATGTAAATGGTTCTTTGGTGTTTTTCAGAAAGAATGTGATGAAGTGAAAAATTTGCATCACAAACCTGAAAAGAAATCCGTTTCCGGTTGCAAGCTCTTTATTCTCCAAAGCCTTGCGTAGCCTGAAGGTAATCGTTTGCGTAAATTCCTTTGTCAAGACCTGTTATCGCACAATGATTCGGTTATACTGTTCGCCGTTGTCCAACAGGACCGCCTATAAAGGCCAAAAATTTTATTGTTAGCTGAGTCAGGAGATGCGG ccacaacggtttccctctagaaataattttgtttaactttaagaaggagatatacat ATGAGCAAAGGTGAAGAACTGTTTACCGGCGTTGTGCCGATTCTGGTGGAACTGGATGGCGATGTGAACGGTCACAAATTCAGCGTGCGTGGTGAAGGTGAAGGCGATGCCACGATTGGCAAACTGACGCTGAAATTTATCTGCACCACCGGCAAACTGCCGGTGCCGTGGCCGACGCTGGTGACCACCCTGACCTATGGCGTTCAGTGTTTTAGTCGCTATCCGGATCACATGAAACGTCACGATTTCTTTAAATCTGCAATGCCGGAAGGCTATGTGCAGGAACGTACGATTAGCTTTAAAGATGATGGCAAATATAAAACGCGCGCCGTTGTGAAATTTGAAGGCGATACCCTGGTGAACCGCATTGAACTGAAAGGCACGGATTTTAAAGAAGATGGCAATATCCTGGGCCATAAACTGGAATACAACTTTAATAGCCATAATGTTTATATTACGGCGGATAAACAGAAAAATGGCATCAAAGCGAATTTTACCGTTCGCCATAACGTTGAAGATGGCAGTGTGCAGCTGGCAGATCATTATCAGCAGAATACCCCGATTGGTGATGGTCCGGTGCTGCTGCCGGATAATCATTATCTGAGCACGCAGACCGTTCTGTCTAAAGATCCGAACGAAAAAGGCACGCGGGACCACATGGTTCTGCACGAATATGTGAATGCGGCAGGTATTACGTGGAGCCATCCGCAGTTCGAAAAATAAgtcgaccggctgctaacaaagcccgaaaggaagctgagttggctgctgccaccgctgagcaataactagcataaccccttggggcctctaaacgggtcttgaggggttttttgctgaaagccaattctgattagaaaaactcatcgagcatcaaatgaaactgcaatttattcatatcaggattatcaataccatatttttgaaaaagccgtttctgtaatgaaggagaaaactcaccgaggcagttccataggatggcaagatcctggtatcggtctgcgattccgactcgtccaacatcaatacaacctattaatttcccctcgtcaaaaataaggttatcaagtgagaaatcaccatgagtgacgactgaatccggtgagaatggcaaaagcttatgcatttctttccagacttgttcaacaggccagccattacgctcgtcatcaaaatcactcgcatcaaccaaaccgttattcattcgtgattgcgcctgagcgagacgaaatacgcgatcgctgttaaaaggacaattacaaacaggaatcgaatgcaaccggcgcaggaacactgccagcgcatcaacaatattttcacctgaatcaggatattcttctaatacctggaatgctgttttcccggggatcgcagtggtgagtaaccatgcatcatcaggagtacggataaaatgcttgatggtcggaagaggcataaattccgtcagccagtttagtctgaccatctcatctgtaacatcattggcaacgctacctttgccatgtttcagaaacaactctggcgcatcgggcttcccatacaatcgatagattgtcgcacctgattgcccgacattatcgcgagcccatttatacccatataaatcagcatccatgttggaatttaatcgcggcttcgagcaagacgtttcccgttgaatatggctcataacaccccttgtattactgtttatgtaagcagacagttttattgttcatgatgatatatttttatcttgtgcaatgtaacatcagagattttgagacacaacgtg | |

| pEutSsfGFP | Plasmid encoding sfGFP expression under P_EutS_ |
| --- | --- |
| P_EutS_ Stability hairpin RBS sfGFP Terminator Kanamycin resistance cassette ColE1 origin  agatcaaaggatcttcttgagatcctttttttctgcgcgtaatctgctgcttgcaaacaaaaaaaccaccgctaccagcggtggtttgtttgccggatcaagagctaccaactctttttccgaaggtaactggcttcagcagagcgcagataccaaatactgttcttctagtgtagccgtagttaggccaccacttcaagaactctgtagcaccgcctacatacctcgctctgctaatcctgttaccagtggctgctgccagtggcgataagtcgtgtcttaccgggttggactcaagacgatagttaccggataaggcgcagcggtcgggctgaacggggggttcgtgcacacagcccagcttggagcgaacgacctacaccgaactgagatacctacagcgtgagctatgagaaagcgccacgcttcccgaagggagaaaggcggacaggtatccggtaagcggcagggtcggaacaggagagcgcacgagggagcttccagggggaaacgcctggtatctttatagtcctgtcgggtttcgccacctctgacttgagcgtcgatttttgtgatgctcgtcaggggggcggagcctatggaaaaacgccagcaacgcgatcccgcgaaat GCGCTTGCCGAATTTTGTTATTTACTCTGACGAAAAATTGTCACGATACACGAAAGTTTTTCACAGGCGGCGACTC ccacaacggtttccctctagaaataattttgtttaactttaagaaggagatatacat ATGAGCAAAGGTGAAGAACTGTTTACCGGCGTTGTGCCGATTCTGGTGGAACTGGATGGCGATGTGAACGGTCACAAATTCAGCGTGCGTGGTGAAGGTGAAGGCGATGCCACGATTGGCAAACTGACGCTGAAATTTATCTGCACCACCGGCAAACTGCCGGTGCCGTGGCCGACGCTGGTGACCACCCTGACCTATGGCGTTCAGTGTTTTAGTCGCTATCCGGATCACATGAAACGTCACGATTTCTTTAAATCTGCAATGCCGGAAGGCTATGTGCAGGAACGTACGATTAGCTTTAAAGATGATGGCAAATATAAAACGCGCGCCGTTGTGAAATTTGAAGGCGATACCCTGGTGAACCGCATTGAACTGAAAGGCACGGATTTTAAAGAAGATGGCAATATCCTGGGCCATAAACTGGAATACAACTTTAATAGCCATAATGTTTATATTACGGCGGATAAACAGAAAAATGGCATCAAAGCGAATTTTACCGTTCGCCATAACGTTGAAGATGGCAGTGTGCAGCTGGCAGATCATTATCAGCAGAATACCCCGATTGGTGATGGTCCGGTGCTGCTGCCGGATAATCATTATCTGAGCACGCAGACCGTTCTGTCTAAAGATCCGAACGAAAAAGGCACGCGGGACCACATGGTTCTGCACGAATATGTGAATGCGGCAGGTATTACGTGGAGCCATCCGCAGTTCGAAAAATAAgtcgaccggctgctaacaaagcccgaaaggaagctgagttggctgctgccaccgctgagcaataactagcataaccccttggggcctctaaacgggtcttgaggggttttttgctgaaagccaattctgattagaaaaactcatcgagcatcaaatgaaactgcaatttattcatatcaggattatcaataccatatttttgaaaaagccgtttctgtaatgaaggagaaaactcaccgaggcagttccataggatggcaagatcctggtatcggtctgcgattccgactcgtccaacatcaatacaacctattaatttcccctcgtcaaaaataaggttatcaagtgagaaatcaccatgagtgacgactgaatccggtgagaatggcaaaagcttatgcatttctttccagacttgttcaacaggccagccattacgctcgtcatcaaaatcactcgcatcaaccaaaccgttattcattcgtgattgcgcctgagcgagacgaaatacgcgatcgctgttaaaaggacaattacaaacaggaatcgaatgcaaccggcgcaggaacactgccagcgcatcaacaatattttcacctgaatcaggatattcttctaatacctggaatgctgttttcccggggatcgcagtggtgagtaaccatgcatcatcaggagtacggataaaatgcttgatggtcggaagaggcataaattccgtcagccagtttagtctgaccatctcatctgtaacatcattggcaacgctacctttgccatgtttcagaaacaactctggcgcatcgggcttcccatacaatcgatagattgtcgcacctgattgcccgacattatcgcgagcccatttatacccatataaatcagcatccatgttggaatttaatcgcggcttcgagcaagacgtttcccgttgaatatggctcataacaccccttgtattactgtttatgtaagcagacagttttattgttcatgatgatatatttttatcttgtgcaatgtaacatcagagattttgagacacaacgtg | |

| pZntR | Plasmid encoding EutR expression under P_T7_ |
| --- | --- |
| P_T7_ Stability hairpin RBS zntR Terminator Kanamycin resistance cassette ColE1 origin  agatcaaaggatcttcttgagatcctttttttctgcgcgtaatctgctgcttgcaaacaaaaaaaccaccgctaccagcggtggtttgtttgccggatcaagagctaccaactctttttccgaaggtaactggcttcagcagagcgcagataccaaatactgttcttctagtgtagccgtagttaggccaccacttcaagaactctgtagcaccgcctacatacctcgctctgctaatcctgttaccagtggctgctgccagtggcgataagtcgtgtcttaccgggttggactcaagacgatagttaccggataaggcgcagcggtcgggctgaacggggggttcgtgcacacagcccagcttggagcgaacgacctacaccgaactgagatacctacagcgtgagctatgagaaagcgccacgcttcccgaagggagaaaggcggacaggtatccggtaagcggcagggtcggaacaggagagcgcacgagggagcttccagggggaaacgcctggtatctttatagtcctgtcgggtttcgccacctctgacttgagcgtcgatttttgtgatgctcgtcaggggggcggagcctatggaaaaacgccagcaacgcgatcccgcgaaattaatacgactcactatagggagaccacaacggtttccctctagaaataattttgtttaactttaagaaggagatatacat ATGTATCGCATTGGTGAGCTGGCAAAAATGGCGGAAGTAACACCCGACACGATTCGTTATTACGAAAAACAGCAGATGATGGAGCATGAAGTGCGTACTGAAGGTGGGTTTCGCCTATATACCGAAAGCGATCTCCAGCGATTGAAATTTATCCGCCATGCCAGACAACTAGGTTTCAGTCTGGAGTCGATCCGCGAGTTGCTGTCGATCCGCATCGATCCTGAACACCATACCTGTCAGGAGTCAAAAGGCATTGTGCAGGAAAGATTGCAGGAAGTCGAAGCACGGATAGCCGAGTTGCAGAGTATGCAGCGTTCCTTGCAACGCCTTAACGATGCCTGTTGTGGGACTGCTCATAGCAGTGTTTATTGTTCGATTCTTGAAGCTCTTGAACAAGGGGCGAGTGGCGTTAAGAGTGGTTGTTGATAAgtcgaccggctgctaacaaagcccgaaaggaagctgagttggctgctgccaccgctgagcaataactagcataaccccttggggcctctaaacgggtcttgaggggttttttgctgaaagccaattctgattagaaaaactcatcgagcatcaaatgaaactgcaatttattcatatcaggattatcaataccatatttttgaaaaagccgtttctgtaatgaaggagaaaactcaccgaggcagttccataggatggcaagatcctggtatcggtctgcgattccgactcgtccaacatcaatacaacctattaatttcccctcgtcaaaaataaggttatcaagtgagaaatcaccatgagtgacgactgaatccggtgagaatggcaaaagcttatgcatttctttccagacttgttcaacaggccagccattacgctcgtcatcaaaatcactcgcatcaaccaaaccgttattcattcgtgattgcgcctgagcgagacgaaatacgcgatcgctgttaaaaggacaattacaaacaggaatcgaatgcaaccggcgcaggaacactgccagcgcatcaacaatattttcacctgaatcaggatattcttctaatacctggaatgctgttttcccggggatcgcagtggtgagtaaccatgcatcatcaggagtacggataaaatgcttgatggtcggaagaggcataaattccgtcagccagtttagtctgaccatctcatctgtaacatcattggcaacgctacctttgccatgtttcagaaacaactctggcgcatcgggcttcccatacaatcgatagattgtcgcacctgattgcccgacattatcgcgagcccatttatacccatataaatcagcatccatgttggaatttaatcgcggcttcgagcaagacgtttcccgttgaatatggctcataacaccccttgtattactgtttatgtaagcagacagttttattgttcatgatgatatatttttatcttgtgcaatgtaacatcagagattttgagacacaacgtg | |

| pZntALacZ | Plasmid encoding EutR expression under P_T7_ |
| --- | --- |
| P_ZntA_ Stability hairpin RBS LacZ Terminator Kanamycin resistance cassette ColE1 origin  agatcaaaggatcttcttgagatcctttttttctgcgcgtaatctgctgcttgcaaacaaaaaaaccaccgctaccagcggtggtttgtttgccggatcaagagctaccaactctttttccgaaggtaactggcttcagcagagcgcagataccaaatactgttcttctagtgtagccgtagttaggccaccacttcaagaactctgtagcaccgcctacatacctcgctctgctaatcctgttaccagtggctgctgccagtggcgataagtcgtgtcttaccgggttggactcaagacgatagttaccggataaggcgcagcggtcgggctgaacggggggttcgtgcacacagcccagcttggagcgaacgacctacaccgaactgagatacctacagcgtgagctatgagaaagcgccacgcttcccgaagggagaaaggcggacaggtatccggtaagcggcagggtcggaacaggagagcgcacgagggagcttccagggggaaacgcctggtatctttatagtcctgtcgggtttcgccacctctgacttgagcgtcgatttttgtgatgctcgtcaggggggcggagcctatggaaaaacgccagcaacgcgatcccgcgaaatCTGTATCTCTGATAAAACTTGACTCTGGAGTCGACTCCAGAGTGTATCCTTCGGTTAATccacaacggtttccctctagaaataattttgtttaactttaagaaggagatatacat atgACCATGATTACGGATTCACTGGCCGTCGTTTTACAACGTCGTGACTGGGAAAACCCTGGCGTTACCCAACTTAATCGCCTTGCAGCACATCCCCCTTTCGCCAGCTGGCGTAATAGCGAAGAGGCCCGCACCGATCGCCCTTCCCAACAGTTGCGCAGCCTGAATGGCGAATGGCGCTTTGCCTGGTTTCCGGCACCAGAAGCGGTGCCGGAAAGCTGGCTGGAGTGCGATCTTCCTGAGGCCGATACTGTCGTCGTCCCCTCAAACTGGCAGATGCACGGTTACGATGCGCCCATCTACACCAACGTGACCTATCCCATTACGGTCAATCCGCCGTTTGTTCCCACGGAGAATCCGACGGGTTGTTACTCGCTCACATTTAATGTTGATGAAAGCTGGCTACAGGAAGGCCAGACGCGAATTATTTTTGATGGCGTTAACTCGGCGTTTCATCTGTGGTGCAACGGGCGCTGGGTCGGTTACGGCCAGGACAGTCGTTTGCCGTCTGAATTTGACCTGAGCGCATTTTTACGCGCCGGAGAAAACCGCCTCGCGGTGATGGTGCTGCGCTGGAGTGACGGCAGTTATCTGGAAGATCAGGATATGTGGCGGATGAGCGGCATTTTCCGTGACGTCTCGTTGCTGCATAAACCGACTACACAAATCAGCGATTTCCATGTTGCCACTCGCTTTAATGATGATTTCAGCCGCGCTGTACTGGAGGCTGAAGTTCAGATGTGCGGCGAGTTGCGTGACTACCTACGGGTAACAGTTTCTTTATGGCAGGGTGAAACGCAGGTCGCCAGCGGCACCGCGCCTTTCGGCGGTGAAATTATCGATGAGCGTGGTGGTTATGCCGATCGCGTCACACTACGTCTGAACGTCGAAAACCCGAAACTGTGGAGCGCCGAAATCCCGAATCTCTATCGTGCGGTGGTTGAACTGCACACCGCCGACGGCACGCTGATTGAAGCAGAAGCCTGCGATGTCGGTTTCCGCGAGGTGCGGATTGAAAATGGTCTGCTGCTGCTGAACGGCAAGCCGTTGCTGATTCGAGGCGTTAACCGTCACGAGCATCATCCTCTGCATGGTCAGGTCATGGATGAGCAGACGATGGTGCAGGATATCCTGCTGATGAAGCAGAACAACTTTAACGCCGTGCGCTGTTCGCATTATCCGAACCATCCGCTGTGGTACACGCTGTGCGACCGCTACGGCCTGTATGTGGTGGATGAAGCCAATATTGAAACCCACGGCATGGTGCCAATGAATCGTCTGACCGATGATCCGCGCTGGCTACCGGCGATGAGCGAACGCGTAACGCGAATGGTGCAGCGCGATCGTAATCACCCGAGTGTGATCATCTGGTCGCTGGGGAATGAATCAGGCCACGGCGCTAATCACGACGCGCTGTATCGCTGGATCAAATCTGTCGATCCTTCCCGCCCGGTGCAGTATGAAGGCGGCGGAGCCGACACCACGGCCACCGATATTATTTGCCCGATGTACGCGCGCGTGGATGAAGACCAGCCCTTCCCGGCTGTGCCGAAATGGTCCATCAAAAAATGGCTTTCGCTACCTGGAGAGACGCGCCCGCTGATCCTTTGCGAATACGCCCACGCGATGGGTAACAGTCTTGGCGGTTTCGCTAAATACTGGCAGGCGTTTCGTCAGTATCCCCGTTTACAGGGCGGCTTCGTCTGGGACTGGGTGGATCAGTCGCTGATTAAATATGATGAAAACGGCAACCCGTGGTCGGCTTACGGCGGTGATTTTGGCGATACGCCGAACGATCGCCAGTTCTGTATGAACGGTCTGGTCTTTGCCGACCGCACGCCGCATCCAGCGCTGACGGAAGCAAAACACCAGCAGCAGTTTTTCCAGTTCCGTTTATCCGGGCAAACCATCGAAGTGACCAGCGAATACCTGTTCCGTCATAGCGATAACGAGCTCCTGCACTGGATGGTGGCGCTGGATGGTAAGCCGCTGGCAAGCGGTGAAGTGCCTCTGGATGTCGCTCCACAAGGTAAACAGTTGATTGAACTGCCTGAACTACCGCAGCCGGAGAGCGCCGGGCAACTCTGGCTCACAGTACGCGTAGTGCAACCGAACGCGACCGCATGGTCAGAAGCCGGACACATCAGCGCCTGGCAGCAGTGGCGTCTGGCTGAAAACCTCAGCGTGACACTCCCCGCCGCGTCCCACGCCATCCCGCATCTGACCACCAGCGAAATGGATTTTTGCATCGAGCTGGGTAATAAGCGTTGGCAATTTAACCGCCAGTCAGGCTTTCTTTCACAGATGTGGATTGGCGATAAAAAACAACTGCTGACGCCGCTGCGCGATCAGTTCACCCGTGCACCGCTGGATAACGACATTGGCGTAAGTGAAGCGACCCGCATTGACCCTAACGCCTGGGTCGAACGCTGGAAGGCGGCGGGCCATTACCAGGCCGAAGCAGCGTTGTTGCAGTGCACGGCAGATACACTTGCTGATGCGGTGCTGATTACGACCGCTCACGCGTGGCAGCATCAGGGGAAAACCTTATTTATCAGCCGGAAAACCTACCGGATTGATGGTAGTGGTCAAATGGCGATTACCGTTGATGTTGAAGTGGCGAGCGATACACCGCATCCGGCGCGGATTGGCCTGAACTGCCAGCTGGCGCAGGTAGCAGAGCGGGTAAACTGGCTCGGATTAGGGCCGCAAGAAAACTATCCCGACCGCCTTACTGCCGCCTGTTTTGACCGCTGGGATCTGCCATTGTCAGACATGTATACCCCGTACGTCTTCCCGAGCGAAAACGGTCTGCGCTGCGGGACGCGCGAATTGAATTATGGCCCACACCAGTGGCGCGGCGACTTCCAGTTCAACATCAGCCGCTACAGTCAACAGCAACTGATGGAAACCAGCCATCGCCATCTGCTGCACGCGGAAGAAGGCACATGGCTGAATATCGACGGTTTCCATATGGGGATTGGTGGCGACGACTCCTGGAGCCCGTCAGTATCGGCGGAATTCCAGCTGAGCGCCGGTCGCTACCATTACCAGTTGGTCTGGTGTCAAAAAtaagtcgaccggctgctaacaaagcccgaaaggaagctgagttggctgctgccaccgctgagcaataactagcataaccccttggggcctctaaacgggtcttgaggggttttttgctgaaagccaattctgattagaaaaactcatcgagcatcaaatgaaactgcaatttattcatatcaggattatcaataccatatttttgaaaaagccgtttctgtaatgaaggagaaaactcaccgaggcagttccataggatggcaagatcctggtatcggtctgcgattccgactcgtccaacatcaatacaacctattaatttcccctcgtcaaaaataaggttatcaagtgagaaatcaccatgagtgacgactgaatccggtgagaatggcaaaagcttatgcatttctttccagacttgttcaacaggccagccattacgctcgtcatcaaaatcactcgcatcaaccaaaccgttattcattcgtgattgcgcctgagcgagacgaaatacgcgatcgctgttaaaaggacaattacaaacaggaatcgaatgcaaccggcgcaggaacactgccagcgcatcaacaatattttcacctgaatcaggatattcttctaatacctggaatgctgttttcccggggatcgcagtggtgagtaaccatgcatcatcaggagtacggataaaatgcttgatggtcggaagaggcataaattccgtcagccagtttagtctgaccatctcatctgtaacatcattggcaacgctacctttgccatgtttcagaaacaactctggcgcatcgggcttcccatacaatcgatagattgtcgcacctgattgcccgacattatcgcgagcccatttatacccatataaatcagcatccatgttggaatttaatcgcggcttcgagcaagacgtttcccgttgaatatggctcataacaccccttgtattactgtttatgtaagcagacagttttattgttcatgatgatatatttttatcttgtgcaatgtaacatcagagattttgagacacaacgtg | |

| P_T7 sfGFP_ | Plasmid encoding sfGFP expression under P_T7_ |
| --- | --- |
| P_T7_ Stability hairpin RBS sfGFP Terminator Kanamycin resistance cassette ColE1 origin    agatcaaaggatcttcttgagatcctttttttctgcgcgtaatctgctgcttgcaaacaaaaaaaccaccgctaccagcggtggtttgtttgccggatcaagagctaccaactctttttccgaaggtaactggcttcagcagagcgcagataccaaatactgttcttctagtgtagccgtagttaggccaccacttcaagaactctgtagcaccgcctacatacctcgctctgctaatcctgttaccagtggctgctgccagtggcgataagtcgtgtcttaccgggttggactcaagacgatagttaccggataaggcgcagcggtcgggctgaacggggggttcgtgcacacagcccagcttggagcgaacgacctacaccgaactgagatacctacagcgtgagctatgagaaagcgccacgcttcccgaagggagaaaggcggacaggtatccggtaagcggcagggtcggaacaggagagcgcacgagggagcttccagggggaaacgcctggtatctttatagtcctgtcgggtttcgccacctctgacttgagcgtcgatttttgtgatgctcgtcaggggggcggagcctatggaaaaacgccagcaacgcgatcccgcgaaattaatacgactcactatagggagaccacaacggtttccctctagaaataattttgtttaactttaagaaggagatatacatATGAGCAAAGGTGAAGAACTGTTTACCGGCGTTGTGCCGATTCTGGTGGAACTGGATGGCGATGTGAACGGTCACAAATTCAGCGTGCGTGGTGAAGGTGAAGGCGATGCCACGATTGGCAAACTGACGCTGAAATTTATCTGCACCACCGGCAAACTGCCGGTGCCGTGGCCGACGCTGGTGACCACCCTGACCTATGGCGTTCAGTGTTTTAGTCGCTATCCGGATCACATGAAACGTCACGATTTCTTTAAATCTGCAATGCCGGAAGGCTATGTGCAGGAACGTACGATTAGCTTTAAAGATGATGGCAAATATAAAACGCGCGCCGTTGTGAAATTTGAAGGCGATACCCTGGTGAACCGCATTGAACTGAAAGGCACGGATTTTAAAGAAGATGGCAATATCCTGGGCCATAAACTGGAATACAACTTTAATAGCCATAATGTTTATATTACGGCGGATAAACAGAAAAATGGCATCAAAGCGAATTTTACCGTTCGCCATAACGTTGAAGATGGCAGTGTGCAGCTGGCAGATCATTATCAGCAGAATACCCCGATTGGTGATGGTCCGGTGCTGCTGCCGGATAATCATTATCTGAGCACGCAGACCGTTCTGTCTAAAGATCCGAACGAAAAAGGCACGCGGGACCACATGGTTCTGCACGAATATGTGAATGCGGCAGGTATTACGTGGAGCCATCCGCAGTTCGAAAAATAAgtcgaccggctgctaacaaagcccgaaaggaagctgagttggctgctgccaccgctgagcaataactagcataaccccttggggcctctaaacgggtcttgaggggttttttgctgaaagccaattctgattagaaaaactcatcgagcatcaaatgaaactgcaatttattcatatcaggattatcaataccatatttttgaaaaagccgtttctgtaatgaaggagaaaactcaccgaggcagttccataggatggcaagatcctggtatcggtctgcgattccgactcgtccaacatcaatacaacctattaatttcccctcgtcaaaaataaggttatcaagtgagaaatcaccatgagtgacgactgaatccggtgagaatggcaaaagcttatgcatttctttccagacttgttcaacaggccagccattacgctcgtcatcaaaatcactcgcatcaaccaaaccgttattcattcgtgattgcgcctgagcgagacgaaatacgcgatcgctgttaaaaggacaattacaaacaggaatcgaatgcaaccggcgcaggaacactgccagcgcatcaacaatattttcacctgaatcaggatattcttctaatacctggaatgctgttttcccggggatcgcagtggtgagtaaccatgcatcatcaggagtacggataaaatgcttgatggtcggaagaggcataaattccgtcagccagtttagtctgaccatctcatctgtaacatcattggcaacgctacctttgccatgtttcagaaacaactctggcgcatcgggcttcccatacaatcgatagattgtcgcacctgattgcccgacattatcgcgagcccatttatacccatataaatcagcatccatgttggaatttaatcgcggcttcgagcaagacgtttcccgttgaatatggctcataacaccccttgtattactgtttatgtaagcagacagttttattgttcatgatgatatatttttatcttgtgcaatgtaacatcagagattttgagacacaacgtg | |

| P_T7,73 sfGFP_ | Plasmid encoding sfGFP expression under P_T7,73_ |
| --- | --- |
| P_T7,73_ Stability hairpin RBS sfGFP Terminator Kanamycin resistance cassette ColE1 origin    agatcaaaggatcttcttgagatcctttttttctgcgcgtaatctgctgcttgcaaacaaaaaaaccaccgctaccagcggtggtttgtttgccggatcaagagctaccaactctttttccgaaggtaactggcttcagcagagcgcagataccaaatactgttcttctagtgtagccgtagttaggccaccacttcaagaactctgtagcaccgcctacatacctcgctctgctaatcctgttaccagtggctgctgccagtggcgataagtcgtgtcttaccgggttggactcaagacgatagttaccggataaggcgcagcggtcgggctgaacggggggttcgtgcacacagcccagcttggagcgaacgacctacaccgaactgagatacctacagcgtgagctatgagaaagcgccacgcttcccgaagggagaaaggcggacaggtatccggtaagcggcagggtcggaacaggagagcgcacgagggagcttccagggggaaacgcctggtatctttatagtcctgtcgggtttcgccacctctgacttgagcgtcgatttttgtgatgctcgtcaggggggcggagcctatggaaaaacgccagcaacgcgatcccgcgaaatTAATACGACTCACTAAAGGgagaccacaacggtttccctctagaaataattttgtttaactttaagaaggagatatacatATGAGCAAAGGTGAAGAACTGTTTACCGGCGTTGTGCCGATTCTGGTGGAACTGGATGGCGATGTGAACGGTCACAAATTCAGCGTGCGTGGTGAAGGTGAAGGCGATGCCACGATTGGCAAACTGACGCTGAAATTTATCTGCACCACCGGCAAACTGCCGGTGCCGTGGCCGACGCTGGTGACCACCCTGACCTATGGCGTTCAGTGTTTTAGTCGCTATCCGGATCACATGAAACGTCACGATTTCTTTAAATCTGCAATGCCGGAAGGCTATGTGCAGGAACGTACGATTAGCTTTAAAGATGATGGCAAATATAAAACGCGCGCCGTTGTGAAATTTGAAGGCGATACCCTGGTGAACCGCATTGAACTGAAAGGCACGGATTTTAAAGAAGATGGCAATATCCTGGGCCATAAACTGGAATACAACTTTAATAGCCATAATGTTTATATTACGGCGGATAAACAGAAAAATGGCATCAAAGCGAATTTTACCGTTCGCCATAACGTTGAAGATGGCAGTGTGCAGCTGGCAGATCATTATCAGCAGAATACCCCGATTGGTGATGGTCCGGTGCTGCTGCCGGATAATCATTATCTGAGCACGCAGACCGTTCTGTCTAAAGATCCGAACGAAAAAGGCACGCGGGACCACATGGTTCTGCACGAATATGTGAATGCGGCAGGTATTACGTGGAGCCATCCGCAGTTCGAAAAATAAgtcgaccggctgctaacaaagcccgaaaggaagctgagttggctgctgccaccgctgagcaataactagcataaccccttggggcctctaaacgggtcttgaggggttttttgctgaaagccaattctgattagaaaaactcatcgagcatcaaatgaaactgcaatttattcatatcaggattatcaataccatatttttgaaaaagccgtttctgtaatgaaggagaaaactcaccgaggcagttccataggatggcaagatcctggtatcggtctgcgattccgactcgtccaacatcaatacaacctattaatttcccctcgtcaaaaataaggttatcaagtgagaaatcaccatgagtgacgactgaatccggtgagaatggcaaaagcttatgcatttctttccagacttgttcaacaggccagccattacgctcgtcatcaaaatcactcgcatcaaccaaaccgttattcattcgtgattgcgcctgagcgagacgaaatacgcgatcgctgttaaaaggacaattacaaacaggaatcgaatgcaaccggcgcaggaacactgccagcgcatcaacaatattttcacctgaatcaggatattcttctaatacctggaatgctgttttcccggggatcgcagtggtgagtaaccatgcatcatcaggagtacggataaaatgcttgatggtcggaagaggcataaattccgtcagccagtttagtctgaccatctcatctgtaacatcattggcaacgctacctttgccatgtttcagaaacaactctggcgcatcgggcttcccatacaatcgatagattgtcgcacctgattgcccgacattatcgcgagcccatttatacccatataaatcagcatccatgttggaatttaatcgcggcttcgagcaagacgtttcccgttgaatatggctcataacaccccttgtattactgtttatgtaagcagacagttttattgttcatgatgatatatttttatcttgtgcaatgtaacatcagagattttgagacacaacgtg | |

| P _σ70,J23119 sfGFP_ | Plasmid encoding sfGFP expression under P _J23119_ |
| --- | --- |
| P _J23119_ Stability hairpin RBS sfGFP Terminator Kanamycin resistance cassette ColE1 origin    agatcaaaggatcttcttgagatcctttttttctgcgcgtaatctgctgcttgcaaacaaaaaaaccaccgctaccagcggtggtttgtttgccggatcaagagctaccaactctttttccgaaggtaactggcttcagcagagcgcagataccaaatactgttcttctagtgtagccgtagttaggccaccacttcaagaactctgtagcaccgcctacatacctcgctctgctaatcctgttaccagtggctgctgccagtggcgataagtcgtgtcttaccgggttggactcaagacgatagttaccggataaggcgcagcggtcgggctgaacggggggttcgtgcacacagcccagcttggagcgaacgacctacaccgaactgagatacctacagcgtgagctatgagaaagcgccacgcttcccgaagggagaaaggcggacaggtatccggtaagcggcagggtcggaacaggagagcgcacgagggagcttccagggggaaacgcctggtatctttatagtcctgtcgggtttcgccacctctgacttgagcgtcgatttttgtgatgctcgtcaggggggcggagcctatggaaaaacgccagcaacgcgatcccgcgaaatttgacagctagctcagtcctaggtataatactagtccacaacggtttccctctagaaataattttgtttaactttaagaaggagatatacatATGAGCAAAGGTGAAGAACTGTTTACCGGCGTTGTGCCGATTCTGGTGGAACTGGATGGCGATGTGAACGGTCACAAATTCAGCGTGCGTGGTGAAGGTGAAGGCGATGCCACGATTGGCAAACTGACGCTGAAATTTATCTGCACCACCGGCAAACTGCCGGTGCCGTGGCCGACGCTGGTGACCACCCTGACCTATGGCGTTCAGTGTTTTAGTCGCTATCCGGATCACATGAAACGTCACGATTTCTTTAAATCTGCAATGCCGGAAGGCTATGTGCAGGAACGTACGATTAGCTTTAAAGATGATGGCAAATATAAAACGCGCGCCGTTGTGAAATTTGAAGGCGATACCCTGGTGAACCGCATTGAACTGAAAGGCACGGATTTTAAAGAAGATGGCAATATCCTGGGCCATAAACTGGAATACAACTTTAATAGCCATAATGTTTATATTACGGCGGATAAACAGAAAAATGGCATCAAAGCGAATTTTACCGTTCGCCATAACGTTGAAGATGGCAGTGTGCAGCTGGCAGATCATTATCAGCAGAATACCCCGATTGGTGATGGTCCGGTGCTGCTGCCGGATAATCATTATCTGAGCACGCAGACCGTTCTGTCTAAAGATCCGAACGAAAAAGGCACGCGGGACCACATGGTTCTGCACGAATATGTGAATGCGGCAGGTATTACGTGGAGCCATCCGCAGTTCGAAAAATAAgtcgaccggctgctaacaaagcccgaaaggaagctgagttggctgctgccagaagcttgggcccgaacaaaaactcatctcagaagaggatctgaatagcgccgtcgaccatcatcatcatcatcattgagtttaaacggtctccagcttggctgttttggcggatgagagaagattttcagcctgatacagattaaatcagaacgcagaagcggtctgataaaacagaatttgcctggcggcagtagcgcggtggtcccacctgaccccatgccgaactcagaagtgaaacgccgtagcgccgatggtagtgtggggtctccccatgcgagagtagggaactgccaggcatcaaataaaacgaaaggctcagtcgaaagactgggcctttcgttttatctgttgtttgtcggtgaactctgaaagccaattctgattagaaaaactcatcgagcatcaaatgaaactgcaatttattcatatcaggattatcaataccatatttttgaaaaagccgtttctgtaatgaaggagaaaactcaccgaggcagttccataggatggcaagatcctggtatcggtctgcgattccgactcgtccaacatcaatacaacctattaatttcccctcgtcaaaaataaggttatcaagtgagaaatcaccatgagtgacgactgaatccggtgagaatggcaaaagcttatgcatttctttccagacttgttcaacaggccagccattacgctcgtcatcaaaatcactcgcatcaaccaaaccgttattcattcgtgattgcgcctgagcgagacgaaatacgcgatcgctgttaaaaggacaattacaaacaggaatcgaatgcaaccggcgcaggaacactgccagcgcatcaacaatattttcacctgaatcaggatattcttctaatacctggaatgctgttttcccggggatcgcagtggtgagtaaccatgcatcatcaggagtacggataaaatgcttgatggtcggaagaggcataaattccgtcagccagtttagtctgaccatctcatctgtaacatcattggcaacgctacctttgccatgtttcagaaacaactctggcgcatcgggcttcccatacaatcgatagattgtcgcacctgattgcccgacattatcgcgagcccatttatacccatataaatcagcatccatgttggaatttaatcgcggcttcgagcaagacgtttcccgttgaatatggctcataacaccccttgtattactgtttatgtaagcagacagttttattgttcatgatgatatatttttatcttgtgcaatgtaacatcagagattttgagacacaacgtg | |

| P_σ70,J23105 sfGFP_ | Plasmid encoding sfGFP expression under P_J23105_ |
| --- | --- |
| P _J23105_Stability hairpin RBS sfGFP Terminator Kanamycin resistance cassette ColE1 origin    agatcaaaggatcttcttgagatcctttttttctgcgcgtaatctgctgcttgcaaacaaaaaaaccaccgctaccagcggtggtttgtttgccggatcaagagctaccaactctttttccgaaggtaactggcttcagcagagcgcagataccaaatactgttcttctagtgtagccgtagttaggccaccacttcaagaactctgtagcaccgcctacatacctcgctctgctaatcctgttaccagtggctgctgccagtggcgataagtcgtgtcttaccgggttggactcaagacgatagttaccggataaggcgcagcggtcgggctgaacggggggttcgtgcacacagcccagcttggagcgaacgacctacaccgaactgagatacctacagcgtgagctatgagaaagcgccacgcttcccgaagggagaaaggcggacaggtatccggtaagcggcagggtcggaacaggagagcgcacgagggagcttccagggggaaacgcctggtatctttatagtcctgtcgggtttcgccacctctgacttgagcgtcgatttttgtgatgctcgtcaggggggcggagcctatggaaaaacgccagcaacgcgatcccgcgaaattttacggctagctcagtcctaggtactatgctagcccacaacggtttccctctagaaataattttgtttaactttaagaaggagatatacatATGAGCAAAGGTGAAGAACTGTTTACCGGCGTTGTGCCGATTCTGGTGGAACTGGATGGCGATGTGAACGGTCACAAATTCAGCGTGCGTGGTGAAGGTGAAGGCGATGCCACGATTGGCAAACTGACGCTGAAATTTATCTGCACCACCGGCAAACTGCCGGTGCCGTGGCCGACGCTGGTGACCACCCTGACCTATGGCGTTCAGTGTTTTAGTCGCTATCCGGATCACATGAAACGTCACGATTTCTTTAAATCTGCAATGCCGGAAGGCTATGTGCAGGAACGTACGATTAGCTTTAAAGATGATGGCAAATATAAAACGCGCGCCGTTGTGAAATTTGAAGGCGATACCCTGGTGAACCGCATTGAACTGAAAGGCACGGATTTTAAAGAAGATGGCAATATCCTGGGCCATAAACTGGAATACAACTTTAATAGCCATAATGTTTATATTACGGCGGATAAACAGAAAAATGGCATCAAAGCGAATTTTACCGTTCGCCATAACGTTGAAGATGGCAGTGTGCAGCTGGCAGATCATTATCAGCAGAATACCCCGATTGGTGATGGTCCGGTGCTGCTGCCGGATAATCATTATCTGAGCACGCAGACCGTTCTGTCTAAAGATCCGAACGAAAAAGGCACGCGGGACCACATGGTTCTGCACGAATATGTGAATGCGGCAGGTATTACGTGGAGCCATCCGCAGTTCGAAAAATAAgtcgaccggctgctaacaaagcccgaaaggaagctgagttggctgctgccagaagcttgggcccgaacaaaaactcatctcagaagaggatctgaatagcgccgtcgaccatcatcatcatcatcattgagtttaaacggtctccagcttggctgttttggcggatgagagaagattttcagcctgatacagattaaatcagaacgcagaagcggtctgataaaacagaatttgcctggcggcagtagcgcggtggtcccacctgaccccatgccgaactcagaagtgaaacgccgtagcgccgatggtagtgtggggtctccccatgcgagagtagggaactgccaggcatcaaataaaacgaaaggctcagtcgaaagactgggcctttcgttttatctgttgtttgtcggtgaactctgaaagccaattctgattagaaaaactcatcgagcatcaaatgaaactgcaatttattcatatcaggattatcaataccatatttttgaaaaagccgtttctgtaatgaaggagaaaactcaccgaggcagttccataggatggcaagatcctggtatcggtctgcgattccgactcgtccaacatcaatacaacctattaatttcccctcgtcaaaaataaggttatcaagtgagaaatcaccatgagtgacgactgaatccggtgagaatggcaaaagcttatgcatttctttccagacttgttcaacaggccagccattacgctcgtcatcaaaatcactcgcatcaaccaaaccgttattcattcgtgattgcgcctgagcgagacgaaatacgcgatcgctgttaaaaggacaattacaaacaggaatcgaatgcaaccggcgcaggaacactgccagcgcatcaacaatattttcacctgaatcaggatattcttctaatacctggaatgctgttttcccggggatcgcagtggtgagtaaccatgcatcatcaggagtacggataaaatgcttgatggtcggaagaggcataaattccgtcagccagtttagtctgaccatctcatctgtaacatcattggcaacgctacctttgccatgtttcagaaacaactctggcgcatcgggcttcccatacaatcgatagattgtcgcacctgattgcccgacattatcgcgagcccatttatacccatataaatcagcatccatgttggaatttaatcgcggcttcgagcaagacgtttcccgttgaatatggctcataacaccccttgtattactgtttatgtaagcagacagttttattgttcatgatgatatatttttatcttgtgcaatgtaacatcagagattttgagacacaacgtg | |

| P_σ70,J23102 sfGFP_ | Plasmid encoding sfGFP expression under P_J23102_ |
| --- | --- |
| P _J23102_Stability hairpin RBS sfGFP Terminator Kanamycin resistance cassette ColE1 origin    agatcaaaggatcttcttgagatcctttttttctgcgcgtaatctgctgcttgcaaacaaaaaaaccaccgctaccagcggtggtttgtttgccggatcaagagctaccaactctttttccgaaggtaactggcttcagcagagcgcagataccaaatactgttcttctagtgtagccgtagttaggccaccacttcaagaactctgtagcaccgcctacatacctcgctctgctaatcctgttaccagtggctgctgccagtggcgataagtcgtgtcttaccgggttggactcaagacgatagttaccggataaggcgcagcggtcgggctgaacggggggttcgtgcacacagcccagcttggagcgaacgacctacaccgaactgagatacctacagcgtgagctatgagaaagcgccacgcttcccgaagggagaaaggcggacaggtatccggtaagcggcagggtcggaacaggagagcgcacgagggagcttccagggggaaacgcctggtatctttatagtcctgtcgggtttcgccacctctgacttgagcgtcgatttttgtgatgctcgtcaggggggcggagcctatggaaaaacgccagcaacgcgatcccgcgaaatttgacagctagctcagtcctaggtactgtgctagcccacaacggtttccctctagaaataattttgtttaactttaagaaggagatatacatATGAGCAAAGGTGAAGAACTGTTTACCGGCGTTGTGCCGATTCTGGTGGAACTGGATGGCGATGTGAACGGTCACAAATTCAGCGTGCGTGGTGAAGGTGAAGGCGATGCCACGATTGGCAAACTGACGCTGAAATTTATCTGCACCACCGGCAAACTGCCGGTGCCGTGGCCGACGCTGGTGACCACCCTGACCTATGGCGTTCAGTGTTTTAGTCGCTATCCGGATCACATGAAACGTCACGATTTCTTTAAATCTGCAATGCCGGAAGGCTATGTGCAGGAACGTACGATTAGCTTTAAAGATGATGGCAAATATAAAACGCGCGCCGTTGTGAAATTTGAAGGCGATACCCTGGTGAACCGCATTGAACTGAAAGGCACGGATTTTAAAGAAGATGGCAATATCCTGGGCCATAAACTGGAATACAACTTTAATAGCCATAATGTTTATATTACGGCGGATAAACAGAAAAATGGCATCAAAGCGAATTTTACCGTTCGCCATAACGTTGAAGATGGCAGTGTGCAGCTGGCAGATCATTATCAGCAGAATACCCCGATTGGTGATGGTCCGGTGCTGCTGCCGGATAATCATTATCTGAGCACGCAGACCGTTCTGTCTAAAGATCCGAACGAAAAAGGCACGCGGGACCACATGGTTCTGCACGAATATGTGAATGCGGCAGGTATTACGTGGAGCCATCCGCAGTTCGAAAAATAAgtcgaccggctgctaacaaagcccgaaaggaagctgagttggctgctgccagaagcttgggcccgaacaaaaactcatctcagaagaggatctgaatagcgccgtcgaccatcatcatcatcatcattgagtttaaacggtctccagcttggctgttttggcggatgagagaagattttcagcctgatacagattaaatcagaacgcagaagcggtctgataaaacagaatttgcctggcggcagtagcgcggtggtcccacctgaccccatgccgaactcagaagtgaaacgccgtagcgccgatggtagtgtggggtctccccatgcgagagtagggaactgccaggcatcaaataaaacgaaaggctcagtcgaaagactgggcctttcgttttatctgttgtttgtcggtgaactctgaaagccaattctgattagaaaaactcatcgagcatcaaatgaaactgcaatttattcatatcaggattatcaataccatatttttgaaaaagccgtttctgtaatgaaggagaaaactcaccgaggcagttccataggatggcaagatcctggtatcggtctgcgattccgactcgtccaacatcaatacaacctattaatttcccctcgtcaaaaataaggttatcaagtgagaaatcaccatgagtgacgactgaatccggtgagaatggcaaaagcttatgcatttctttccagacttgttcaacaggccagccattacgctcgtcatcaaaatcactcgcatcaaccaaaccgttattcattcgtgattgcgcctgagcgagacgaaatacgcgatcgctgttaaaaggacaattacaaacaggaatcgaatgcaaccggcgcaggaacactgccagcgcatcaacaatattttcacctgaatcaggatattcttctaatacctggaatgctgttttcccggggatcgcagtggtgagtaaccatgcatcatcaggagtacggataaaatgcttgatggtcggaagaggcataaattccgtcagccagtttagtctgaccatctcatctgtaacatcattggcaacgctacctttgccatgtttcagaaacaactctggcgcatcgggcttcccatacaatcgatagattgtcgcacctgattgcccgacattatcgcgagcccatttatacccatataaatcagcatccatgttggaatttaatcgcggcttcgagcaagacgtttcccgttgaatatggctcataacaccccttgtattactgtttatgtaagcagacagttttattgttcatgatgatatatttttatcttgtgcaatgtaacatcagagattttgagacacaacgtg | |

| P_T7 Zur_ | Plasmid encoding Zur expression under P_T7_ |
| --- | --- |
| P_T7_ Stability hairpin RBS Zur Terminator Kanamycin resistance cassette ColE1 origin    agatcaaaggatcttcttgagatcctttttttctgcgcgtaatctgctgcttgcaaacaaaaaaaccaccgctaccagcggtggtttgtttgccggatcaagagctaccaactctttttccgaaggtaactggcttcagcagagcgcagataccaaatactgttcttctagtgtagccgtagttaggccaccacttcaagaactctgtagcaccgcctacatacctcgctctgctaatcctgttaccagtggctgctgccagtggcgataagtcgtgtcttaccgggttggactcaagacgatagttaccggataaggcgcagcggtcgggctgaacggggggttcgtgcacacagcccagcttggagcgaacgacctacaccgaactgagatacctacagcgtgagctatgagaaagcgccacgcttcccgaagggagaaaggcggacaggtatccggtaagcggcagggtcggaacaggagagcgcacgagggagcttccagggggaaacgcctggtatctttatagtcctgtcgggtttcgccacctctgacttgagcgtcgatttttgtgatgctcgtcaggggggcggagcctatggaaaaacgccagcaacgcgatcccgcgaaattaatacgactcactatagggagaccacaacggtttccctctagaaataattttgtttaactttaagaaggagatatacatATGGAAAAGACCACAACGCAGGAGTTATTAGCGCAGGCTGAAAAAATCTGCGCGCAGCGTAATGTGCGCCTGACCCCACAGCGCCTGGAAGTGTTGCGCCTGATGAGTCTCCAAGATGGCGCTATCAGCGCTTATGATCTGCTTGATTTACTGCGCGAAGCTGAACCGCAAGCCAAGCCGCCAACGGTTTATCGCGCGCTGGATTTTCTGCTTGAGCAAGGTTTTGTGCATAAGGTGGAATCCACCAACAGTTATGTGCTCTGTCATCTGTTCGATCAGCCCACCCATACGTCAGCCATGTTTATTTGCGATCGCTGCGGCGCAGTGAAAGAAGAGTGTGCAGAAGGCGTGGAAGACATTATGCATACGCTGGCGGCAAAAATGGGGTTTGCCCTGCGGCATAATGTGATTGAAGCACATGGGCTCTGTGCGGCATGTGTAGAAGTGGAAGCGTGTCGTCATCCTGAACAGTGCCAGCATGATCACTCTGTGCAGGTGAAAAAGAAACCGCGTTAAgtcgaccggctgctaacaaagcccgaaaggaagctgagttggctgctgccaccgctgagcaataactagcataaccccttggggcctctaaacgggtcttgaggggttttttgctgaaagccaattctgattagaaaaactcatcgagcatcaaatgaaactgcaatttattcatatcaggattatcaataccatatttttgaaaaagccgtttctgtaatgaaggagaaaactcaccgaggcagttccataggatggcaagatcctggtatcggtctgcgattccgactcgtccaacatcaatacaacctattaatttcccctcgtcaaaaataaggttatcaagtgagaaatcaccatgagtgacgactgaatccggtgagaatggcaaaagcttatgcatttctttccagacttgttcaacaggccagccattacgctcgtcatcaaaatcactcgcatcaaccaaaccgttattcattcgtgattgcgcctgagcgagacgaaatacgcgatcgctgttaaaaggacaattacaaacaggaatcgaatgcaaccggcgcaggaacactgccagcgcatcaacaatattttcacctgaatcaggatattcttctaatacctggaatgctgttttcccggggatcgcagtggtgagtaaccatgcatcatcaggagtacggataaaatgcttgatggtcggaagaggcataaattccgtcagccagtttagtctgaccatctcatctgtaacatcattggcaacgctacctttgccatgtttcagaaacaactctggcgcatcgggcttcccatacaatcgatagattgtcgcacctgattgcccgacattatcgcgagcccatttatacccatataaatcagcatccatgttggaatttaatcgcggcttcgagcaagacgtttcccgttgaatatggctcataacaccccttgtattactgtttatgtaagcagacagttttattgttcatgatgatatatttttatcttgtgcaatgtaacatcagagattttgagacacaacgtg | |

| P_T7 ZurOsfGFP_ | Plasmid encoding sfGFP expression under ZurO and P_T7_ |
| --- | --- |
| P_T7_ ZurO Stability hairpin RBS sfGFP Terminator Kanamycin resistance cassette ColE1 origin    agatcaaaggatcttcttgagatcctttttttctgcgcgtaatctgctgcttgcaaacaaaaaaaccaccgctaccagcggtggtttgtttgccggatcaagagctaccaactctttttccgaaggtaactggcttcagcagagcgcagataccaaatactgttcttctagtgtagccgtagttaggccaccacttcaagaactctgtagcaccgcctacatacctcgctctgctaatcctgttaccagtggctgctgccagtggcgataagtcgtgtcttaccgggttggactcaagacgatagttaccggataaggcgcagcggtcgggctgaacggggggttcgtgcacacagcccagcttggagcgaacgacctacaccgaactgagatacctacagcgtgagctatgagaaagcgccacgcttcccgaagggagaaaggcggacaggtatccggtaagcggcagggtcggaacaggagagcgcacgagggagcttccagggggaaacgcctggtatctttatagtcctgtcgggtttcgccacctctgacttgagcgtcgatttttgtgatgctcgtcaggggggcggagcctatggaaaaacgccagcaacgcgatcccgcgaaattaatacgactcactatagggagaGAAGTGTGATATTATAACATTTccacaacggtttccctctagaaataattttgtttaactttaagaaggagatatacatATGAGCAAAGGTGAAGAACTGTTTACCGGCGTTGTGCCGATTCTGGTGGAACTGGATGGCGATGTGAACGGTCACAAATTCAGCGTGCGTGGTGAAGGTGAAGGCGATGCCACGATTGGCAAACTGACGCTGAAATTTATCTGCACCACCGGCAAACTGCCGGTGCCGTGGCCGACGCTGGTGACCACCCTGACCTATGGCGTTCAGTGTTTTAGTCGCTATCCGGATCACATGAAACGTCACGATTTCTTTAAATCTGCAATGCCGGAAGGCTATGTGCAGGAACGTACGATTAGCTTTAAAGATGATGGCAAATATAAAACGCGCGCCGTTGTGAAATTTGAAGGCGATACCCTGGTGAACCGCATTGAACTGAAAGGCACGGATTTTAAAGAAGATGGCAATATCCTGGGCCATAAACTGGAATACAACTTTAATAGCCATAATGTTTATATTACGGCGGATAAACAGAAAAATGGCATCAAAGCGAATTTTACCGTTCGCCATAACGTTGAAGATGGCAGTGTGCAGCTGGCAGATCATTATCAGCAGAATACCCCGATTGGTGATGGTCCGGTGCTGCTGCCGGATAATCATTATCTGAGCACGCAGACCGTTCTGTCTAAAGATCCGAACGAAAAAGGCACGCGGGACCACATGGTTCTGCACGAATATGTGAATGCGGCAGGTATTACGTGGAGCCATCCGCAGTTCGAAAAATAAgtcgaccggctgctaacaaagcccgaaaggaagctgagttggctgctgccaccgctgagcaataactagcataaccccttggggcctctaaacgggtcttgaggggttttttgctgaaagccaattctgattagaaaaactcatcgagcatcaaatgaaactgcaatttattcatatcaggattatcaataccatatttttgaaaaagccgtttctgtaatgaaggagaaaactcaccgaggcagttccataggatggcaagatcctggtatcggtctgcgattccgactcgtccaacatcaatacaacctattaatttcccctcgtcaaaaataaggttatcaagtgagaaatcaccatgagtgacgactgaatccggtgagaatggcaaaagcttatgcatttctttccagacttgttcaacaggccagccattacgctcgtcatcaaaatcactcgcatcaaccaaaccgttattcattcgtgattgcgcctgagcgagacgaaatacgcgatcgctgttaaaaggacaattacaaacaggaatcgaatgcaaccggcgcaggaacactgccagcgcatcaacaatattttcacctgaatcaggatattcttctaatacctggaatgctgttttcccggggatcgcagtggtgagtaaccatgcatcatcaggagtacggataaaatgcttgatggtcggaagaggcataaattccgtcagccagtttagtctgaccatctcatctgtaacatcattggcaacgctacctttgccatgtttcagaaacaactctggcgcatcgggcttcccatacaatcgatagattgtcgcacctgattgcccgacattatcgcgagcccatttatacccatataaatcagcatccatgttggaatttaatcgcggcttcgagcaagacgtttcccgttgaatatggctcataacaccccttgtattactgtttatgtaagcagacagttttattgttcatgatgatatatttttatcttgtgcaatgtaacatcagagattttgagacacaacgtg | |

| P_T7 ZurONonReg_ | Plasmid encoding a non-regulatory protein under ZurO and P_T7_ |
| --- | --- |
| P_T7_ ZurO Stability hairpin RBS NonReg Terminator Kanamycin resistance cassette ColE1 origin    agatcaaaggatcttcttgagatcctttttttctgcgcgtaatctgctgcttgcaaacaaaaaaaccaccgctaccagcggtggtttgtttgccggatcaagagctaccaactctttttccgaaggtaactggcttcagcagagcgcagataccaaatactgttcttctagtgtagccgtagttaggccaccacttcaagaactctgtagcaccgcctacatacctcgctctgctaatcctgttaccagtggctgctgccagtggcgataagtcgtgtcttaccgggttggactcaagacgatagttaccggataaggcgcagcggtcgggctgaacggggggttcgtgcacacagcccagcttggagcgaacgacctacaccgaactgagatacctacagcgtgagctatgagaaagcgccacgcttcccgaagggagaaaggcggacaggtatccggtaagcggcagggtcggaacaggagagcgcacgagggagcttccagggggaaacgcctggtatctttatagtcctgtcgggtttcgccacctctgacttgagcgtcgatttttgtgatgctcgtcaggggggcggagcctatggaaaaacgccagcaacgcgatcccgcgaaattaatacgactcactatagggagaGAAGTGTGATATTATAACATTTccacaacggtttccctctagaaataattttgtttaactttaagaaggagatatacat ATGGAAAGAAGGCGTTTCAGCGCCGCCCTGCAAGGGCAGAACCTCAGGTTCACCGGGGCCGGGGAGGGGGGCTCGGCACGGGCGCGGGTGGGCTTCCTGCTGCTCGAACACTTTTCCCTGCCGGCCTTCACCCAGGCGCTGGACACCCTGGTGACCGCCAACCTGATCCGCAGCGAGTCCTTCGCGGTGAGCAACTTCAGCCTGAGCGGCGAGCCGGTGACCAGCGACCTCGGGTTGCTGATCTGCCCGGACCTGGCGCTCGCCTCGGCGGCGTTGCAAGGACTCGACCTGCTGGTGGTCTGCGGCGGTTTGCGCACGCCCCTGCGCTCGTCCGCCGCGCTGCGGGAAGCGTTGCAGGACGTGGCGTCGCGCGGCGTCGCCCTGGCCGGGCTGTGGAACGGTGCCTGGTTCCTCGGCGACGCCGGCCTGCTGGATGGCTACCGCTGCTGCATCCATCCCGAGCACCGCGCCGCCCTCGCCGAGATCGCCCGCAACAGCCAGGTGACCAGCGACAGCTATATGGTCGACCGCGACCGCCTGACCGCGGCCAGCCCCACCGGCGCCTTCAACCTGGCGCTGGAGTGGATCGCCCGGCGCCACGGGCGCGACCTGGTGGAGGCGGTGGTGGATATCCTCGCCTTCGAGGAGTCGCGCTACCGGCGGGTGCGCCCCGCCCTGCACGAGAAGATGAGCGAGCCGCTGCGCGAAGTGATCAACCTGATGTCGGCGAATATCGAGGAACCGCTCAGCCAGGAGCAGCTCGCCCACTACGTGGGGCGCTCGCGGCGGCAGATCGAGCGATTGTTCCAGCAGCAGCTCGGAACCACGCCGGTGCGCTACTACCTGGAGTTGCGGATCACCGAGTGCCGGCGCCTGCTGCAACATTCCGACCTGACCATGCTGGAGGTCCTGGTCGCCTGCGGCTTCGTCTCGCCCAGCCACTTCAGCAAGTGCTACACCGCCTTCTACGGCTACCCGCCGTCGAAAGAAGTGCGCTACGGCAACGTCCGCTCGACGAAATGAgtcgaccggctgctaacaaagcccgaaaggaagctgagttggctgctgccaccgctgagcaataactagcataaccccttggggcctctaaacgggtcttgaggggttttttgctgaaagccaattctgattagaaaaactcatcgagcatcaaatgaaactgcaatttattcatatcaggattatcaataccatatttttgaaaaagccgtttctgtaatgaaggagaaaactcaccgaggcagttccataggatggcaagatcctggtatcggtctgcgattccgactcgtccaacatcaatacaacctattaatttcccctcgtcaaaaataaggttatcaagtgagaaatcaccatgagtgacgactgaatccggtgagaatggcaaaagcttatgcatttctttccagacttgttcaacaggccagccattacgctcgtcatcaaaatcactcgcatcaaccaaaccgttattcattcgtgattgcgcctgagcgagacgaaatacgcgatcgctgttaaaaggacaattacaaacaggaatcgaatgcaaccggcgcaggaacactgccagcgcatcaacaatattttcacctgaatcaggatattcttctaatacctggaatgctgttttcccggggatcgcagtggtgagtaaccatgcatcatcaggagtacggataaaatgcttgatggtcggaagaggcataaattccgtcagccagtttagtctgaccatctcatctgtaacatcattggcaacgctacctttgccatgtttcagaaacaactctggcgcatcgggcttcccatacaatcgatagattgtcgcacctgattgcccgacattatcgcgagcccatttatacccatataaatcagcatccatgttggaatttaatcgcggcttcgagcaagacgtttcccgttgaatatggctcataacaccccttgtattactgtttatgtaagcagacagttttattgttcatgatgatatatttttatcttgtgcaatgtaacatcagagattttgagacacaacgtg | |

| P_T7 ZurONonReg3075_ | Plasmid encoding a non-regulatory protein (3075BPs) under ZurO and P_T7_ |
| --- | --- |
| P_T7_ ZurO Stability hairpin RBS NonReg Terminator Kanamycin resistance cassette ColE1 origin    agatcaaaggatcttcttgagatcctttttttctgcgcgtaatctgctgcttgcaaacaaaaaaaccaccgctaccagcggtggtttgtttgccggatcaagagctaccaactctttttccgaaggtaactggcttcagcagagcgcagataccaaatactgttcttctagtgtagccgtagttaggccaccacttcaagaactctgtagcaccgcctacatacctcgctctgctaatcctgttaccagtggctgctgccagtggcgataagtcgtgtcttaccgggttggactcaagacgatagttaccggataaggcgcagcggtcgggctgaacggggggttcgtgcacacagcccagcttggagcgaacgacctacaccgaactgagatacctacagcgtgagctatgagaaagcgccacgcttcccgaagggagaaaggcggacaggtatccggtaagcggcagggtcggaacaggagagcgcacgagggagcttccagggggaaacgcctggtatctttatagtcctgtcgggtttcgccacctctgacttgagcgtcgatttttgtgatgctcgtcaggggggcggagcctatggaaaaacgccagcaacgcgatcccgcgaaattaatacgactcactatagggagaGAAGTGTGATATTATAACATTTccacaacggtttccctctagaaataattttgtttaactttaagaaggagatatacat atgACCATGATTACGGATTCACTGGCCGTCGTTTTACAACGTCGTGACTGGGAAAACCCTGGCGTTACCCAACTTAATCGCCTTGCAGCACATCCCCCTTTCGCCAGCTGGCGTAATAGCGAAGAGGCCCGCACCGATCGCCCTTCCCAACAGTTGCGCAGCCTGAATGGCGAATGGCGCTTTGCCTGGTTTCCGGCACCAGAAGCGGTGCCGGAAAGCTGGCTGGAGTGCGATCTTCCTGAGGCCGATACTGTCGTCGTCCCCTCAAACTGGCAGATGCACGGTTACGATGCGCCCATCTACACCAACGTGACCTATCCCATTACGGTCAATCCGCCGTTTGTTCCCACGGAGAATCCGACGGGTTGTTACTCGCTCACATTTAATGTTGATGAAAGCTGGCTACAGGAAGGCCAGACGCGAATTATTTTTGATGGCGTTAACTCGGCGTTTCATCTGTGGTGCAACGGGCGCTGGGTCGGTTACGGCCAGGACAGTCGTTTGCCGTCTGAATTTGACCTGAGCGCATTTTTACGCGCCGGAGAAAACCGCCTCGCGGTGATGGTGCTGCGCTGGAGTGACGGCAGTTATCTGGAAGATCAGGATATGTGGCGGATGAGCGGCATTTTCCGTGACGTCTCGTTGCTGCATAAACCGACTACACAAATCAGCGATTTCCATGTTGCCACTCGCTTTAATGATGATTTCAGCCGCGCTGTACTGGAGGCTGAAGTTCAGATGTGCGGCGAGTTGCGTGACTACCTACGGGTAACAGTTTCTTTATGGCAGGGTGAAACGCAGGTCGCCAGCGGCACCGCGCCTTTCGGCGGTGAAATTATCGATGAGCGTGGTGGTTATGCCGATCGCGTCACACTACGTCTGAACGTCGAAAACCCGAAACTGTGGAGCGCCGAAATCCCGAATCTCTATCGTGCGGTGGTTGAACTGCACACCGCCGACGGCACGCTGATTGAAGCAGAAGCCTGCGATGTCGGTTTCCGCGAGGTGCGGATTGAAAATGGTCTGCTGCTGCTGAACGGCAAGCCGTTGCTGATTCGAGGCGTTAACCGTCACGAGCATCATCCTCTGCATGGTCAGGTCATGGATGAGCAGACGATGGTGCAGGATATCCTGCTGATGAAGCAGAACAACTTTAACGCCGTGCGCTGTTCGCATTATCCGAACCATCCGCTGTGGTACACGCTGTGCGACCGCTACGGCCTGTATGTGGTGGATGAAGCCAATATTGAAACCCACGGCATGGTGCCAATGAATCGTCTGACCGATGATCCGCGCTGGCTACCGGCGATGAGCGAACGCGTAACGCGAATGGTGCAGCGCGATCGTAATCACCCGAGTGTGATCATCTGGTCGCTGGGGAATGAATCAGGCCACGGCGCTAATCACGACGCGCTGTATCGCTGGATCAAATCTGTCGATCCTTCCCGCCCGGTGCAGTATGAAGGCGGCGGAGCCGACACCACGGCCACCGATATTATTTGCCCGATGTACGCGCGCGTGGATGAAGACCAGCCCTTCCCGGCTGTGCCGAAATGGTCCATCAAAAAATGGCTTTCGCTACCTGGAGAGACGCGCCCGCTGATCCTTTGCGAATACGCCCACGCGATGGGTAACAGTCTTGGCGGTTTCGCTAAATACTGGCAGGCGTTTCGTCAGTATCCCCGTTTACAGGGCGGCTTCGTCTGGGACTGGGTGGATCAGTCGCTGATTAAATATGATGAAAACGGCAACCCGTGGTCGGCTTACGGCGGTGATTTTGGCGATACGCCGAACGATCGCCAGTTCTGTATGAACGGTCTGGTCTTTGCCGACCGCACGCCGCATCCAGCGCTGACGGAAGCAAAACACCAGCAGCAGTTTTTCCAGTTCCGTTTATCCGGGCAAACCATCGAAGTGACCAGCGAATACCTGTTCCGTCATAGCGATAACGAGCTCCTGCACTGGATGGTGGCGCTGGATGGTAAGCCGCTGGCAAGCGGTGAAGTGCCTCTGGATGTCGCTCCACAAGGTAAACAGTTGATTGAACTGCCTGAACTACCGCAGCCGGAGAGCGCCGGGCAACTCTGGCTCACAGTACGCGTAGTGCAACCGAACGCGACCGCATGGTCAGAAGCCGGACACATCAGCGCCTGGCAGCAGTGGCGTCTGGCTGAAAACCTCAGCGTGACACTCCCCGCCGCGTCCCACGCCATCCCGCATCTGACCACCAGCGAAATGGATTTTTGCATCGAGCTGGGTAATAAGCGTTGGCAATTTAACCGCCAGTCAGGCTTTCTTTCACAGATGTGGATTGGCGATAAAAAACAACTGCTGACGCCGCTGCGCGATCAGTTCACCCGTGCACCGCTGGATAACGACATTGGCGTAAGTGAAGCGACCCGCATTGACCCTAACGCCTGGGTCGAACGCTGGAAGGCGGCGGGCCATTACCAGGCCGAAGCAGCGTTGTTGCAGTGCACGGCAGATACACTTGCTGATGCGGTGCTGATTACGACCGCTCACGCGTGGCAGCATCAGGGGAAAACCTTATTTATCAGCCGGAAAACCTACCGGATTGATGGTAGTGGTCAAATGGCGATTACCGTTGATGTTGAAGTGGCGAGCGATACACCGCATCCGGCGCGGATTGGCCTGAACTGCCAGCTGGCGCAGGTAGCAGAGCGGGTAAACTGGCTCGGATTAGGGCCGCAAGAAAACTATCCCGACCGCCTTACTGCCGCCTGTTTTGACCGCTGGGATCTGCCATTGTCAGACATGTATACCCCGTACGTCTTCCCGAGCGAAAACGGTCTGCGCTGCGGGACGCGCGAATTGAATTATGGCCCACACCAGTGGCGCGGCGACTTCCAGTTCAACATCAGCCGCTACAGTCAACAGCAACTGATGGAAACCAGCCATCGCCATCTGCTGCACGCGGAAGAAGGCACATGGCTGAATATCGACGGTTTCCATATGGGGATTGGTGGCGACGACTCCTGGAGCCCGTCAGTATCGGCGGAATTCCAGCTGAGCGCCGGTCGCTACCATTACCAGTTGGTCTGGTGTCAAAAAtaagtcgaccggctgctaacaaagcccgaaaggaagctgagttggctgctgccaccgctgagcaataactagcataaccccttggggcctctaaacgggtcttgaggggttttttgctgaaagccaattctgattagaaaaactcatcgagcatcaaatgaaactgcaatttattcatatcaggattatcaataccatatttttgaaaaagccgtttctgtaatgaaggagaaaactcaccgaggcagttccataggatggcaagatcctggtatcggtctgcgattccgactcgtccaacatcaatacaacctattaatttcccctcgtcaaaaataaggttatcaagtgagaaatcaccatgagtgacgactgaatccggtgagaatggcaaaagcttatgcatttctttccagacttgttcaacaggccagccattacgctcgtcatcaaaatcactcgcatcaaccaaaccgttattcattcgtgattgcgcctgagcgagacgaaatacgcgatcgctgttaaaaggacaattacaaacaggaatcgaatgcaaccggcgcaggaacactgccagcgcatcaacaatattttcacctgaatcaggatattcttctaatacctggaatgctgttttcccggggatcgcagtggtgagtaaccatgcatcatcaggagtacggataaaatgcttgatggtcggaagaggcataaattccgtcagccagtttagtctgaccatctcatctgtaacatcattggcaacgctacctttgccatgtttcagaaacaactctggcgcatcgggcttcccatacaatcgatagattgtcgcacctgattgcccgacattatcgcgagcccatttatacccatataaatcagcatccatgttggaatttaatcgcggcttcgagcaagacgtttcccgttgaatatggctcataacaccccttgtattactgtttatgtaagcagacagttttattgttcatgatgatatatttttatcttgtgcaatgtaacatcagagattttgagacacaacgtg | |

| P_T7 ZurONonReg3075, NoRBS_ | Plasmid encoding a non-regulatory protein (3075BPs) without an RBS under ZurO and P_T7_ |
| --- | --- |
| P_T7_ ZurO Stability hairpin RBS NonReg Terminator Kanamycin resistance cassette ColE1 origin    agatcaaaggatcttcttgagatcctttttttctgcgcgtaatctgctgcttgcaaacaaaaaaaccaccgctaccagcggtggtttgtttgccggatcaagagctaccaactctttttccgaaggtaactggcttcagcagagcgcagataccaaatactgttcttctagtgtagccgtagttaggccaccacttcaagaactctgtagcaccgcctacatacctcgctctgctaatcctgttaccagtggctgctgccagtggcgataagtcgtgtcttaccgggttggactcaagacgatagttaccggataaggcgcagcggtcgggctgaacggggggttcgtgcacacagcccagcttggagcgaacgacctacaccgaactgagatacctacagcgtgagctatgagaaagcgccacgcttcccgaagggagaaaggcggacaggtatccggtaagcggcagggtcggaacaggagagcgcacgagggagcttccagggggaaacgcctggtatctttatagtcctgtcgggtttcgccacctctgacttgagcgtcgatttttgtgatgctcgtcaggggggcggagcctatggaaaaacgccagcaacgcgatcccgcgaaattaatacgactcactatagggagaGAAGTGTGATATTATAACATTTACCATGATTACGGATTCACTGGCCGTCGTTTTACAACGTCGTGACTGGGAAAACCCTGGCGTTACCCAACTTAATCGCCTTGCAGCACATCCCCCTTTCGCCAGCTGGCGTAATAGCGAAGAGGCCCGCACCGATCGCCCTTCCCAACAGTTGCGCAGCCTGAATGGCGAATGGCGCTTTGCCTGGTTTCCGGCACCAGAAGCGGTGCCGGAAAGCTGGCTGGAGTGCGATCTTCCTGAGGCCGATACTGTCGTCGTCCCCTCAAACTGGCAGATGCACGGTTACGATGCGCCCATCTACACCAACGTGACCTATCCCATTACGGTCAATCCGCCGTTTGTTCCCACGGAGAATCCGACGGGTTGTTACTCGCTCACATTTAATGTTGATGAAAGCTGGCTACAGGAAGGCCAGACGCGAATTATTTTTGATGGCGTTAACTCGGCGTTTCATCTGTGGTGCAACGGGCGCTGGGTCGGTTACGGCCAGGACAGTCGTTTGCCGTCTGAATTTGACCTGAGCGCATTTTTACGCGCCGGAGAAAACCGCCTCGCGGTGATGGTGCTGCGCTGGAGTGACGGCAGTTATCTGGAAGATCAGGATATGTGGCGGATGAGCGGCATTTTCCGTGACGTCTCGTTGCTGCATAAACCGACTACACAAATCAGCGATTTCCATGTTGCCACTCGCTTTAATGATGATTTCAGCCGCGCTGTACTGGAGGCTGAAGTTCAGATGTGCGGCGAGTTGCGTGACTACCTACGGGTAACAGTTTCTTTATGGCAGGGTGAAACGCAGGTCGCCAGCGGCACCGCGCCTTTCGGCGGTGAAATTATCGATGAGCGTGGTGGTTATGCCGATCGCGTCACACTACGTCTGAACGTCGAAAACCCGAAACTGTGGAGCGCCGAAATCCCGAATCTCTATCGTGCGGTGGTTGAACTGCACACCGCCGACGGCACGCTGATTGAAGCAGAAGCCTGCGATGTCGGTTTCCGCGAGGTGCGGATTGAAAATGGTCTGCTGCTGCTGAACGGCAAGCCGTTGCTGATTCGAGGCGTTAACCGTCACGAGCATCATCCTCTGCATGGTCAGGTCATGGATGAGCAGACGATGGTGCAGGATATCCTGCTGATGAAGCAGAACAACTTTAACGCCGTGCGCTGTTCGCATTATCCGAACCATCCGCTGTGGTACACGCTGTGCGACCGCTACGGCCTGTATGTGGTGGATGAAGCCAATATTGAAACCCACGGCATGGTGCCAATGAATCGTCTGACCGATGATCCGCGCTGGCTACCGGCGATGAGCGAACGCGTAACGCGAATGGTGCAGCGCGATCGTAATCACCCGAGTGTGATCATCTGGTCGCTGGGGAATGAATCAGGCCACGGCGCTAATCACGACGCGCTGTATCGCTGGATCAAATCTGTCGATCCTTCCCGCCCGGTGCAGTATGAAGGCGGCGGAGCCGACACCACGGCCACCGATATTATTTGCCCGATGTACGCGCGCGTGGATGAAGACCAGCCCTTCCCGGCTGTGCCGAAATGGTCCATCAAAAAATGGCTTTCGCTACCTGGAGAGACGCGCCCGCTGATCCTTTGCGAATACGCCCACGCGATGGGTAACAGTCTTGGCGGTTTCGCTAAATACTGGCAGGCGTTTCGTCAGTATCCCCGTTTACAGGGCGGCTTCGTCTGGGACTGGGTGGATCAGTCGCTGATTAAATATGATGAAAACGGCAACCCGTGGTCGGCTTACGGCGGTGATTTTGGCGATACGCCGAACGATCGCCAGTTCTGTATGAACGGTCTGGTCTTTGCCGACCGCACGCCGCATCCAGCGCTGACGGAAGCAAAACACCAGCAGCAGTTTTTCCAGTTCCGTTTATCCGGGCAAACCATCGAAGTGACCAGCGAATACCTGTTCCGTCATAGCGATAACGAGCTCCTGCACTGGATGGTGGCGCTGGATGGTAAGCCGCTGGCAAGCGGTGAAGTGCCTCTGGATGTCGCTCCACAAGGTAAACAGTTGATTGAACTGCCTGAACTACCGCAGCCGGAGAGCGCCGGGCAACTCTGGCTCACAGTACGCGTAGTGCAACCGAACGCGACCGCATGGTCAGAAGCCGGACACATCAGCGCCTGGCAGCAGTGGCGTCTGGCTGAAAACCTCAGCGTGACACTCCCCGCCGCGTCCCACGCCATCCCGCATCTGACCACCAGCGAAATGGATTTTTGCATCGAGCTGGGTAATAAGCGTTGGCAATTTAACCGCCAGTCAGGCTTTCTTTCACAGATGTGGATTGGCGATAAAAAACAACTGCTGACGCCGCTGCGCGATCAGTTCACCCGTGCACCGCTGGATAACGACATTGGCGTAAGTGAAGCGACCCGCATTGACCCTAACGCCTGGGTCGAACGCTGGAAGGCGGCGGGCCATTACCAGGCCGAAGCAGCGTTGTTGCAGTGCACGGCAGATACACTTGCTGATGCGGTGCTGATTACGACCGCTCACGCGTGGCAGCATCAGGGGAAAACCTTATTTATCAGCCGGAAAACCTACCGGATTGATGGTAGTGGTCAAATGGCGATTACCGTTGATGTTGAAGTGGCGAGCGATACACCGCATCCGGCGCGGATTGGCCTGAACTGCCAGCTGGCGCAGGTAGCAGAGCGGGTAAACTGGCTCGGATTAGGGCCGCAAGAAAACTATCCCGACCGCCTTACTGCCGCCTGTTTTGACCGCTGGGATCTGCCATTGTCAGACATGTATACCCCGTACGTCTTCCCGAGCGAAAACGGTCTGCGCTGCGGGACGCGCGAATTGAATTATGGCCCACACCAGTGGCGCGGCGACTTCCAGTTCAACATCAGCCGCTACAGTCAACAGCAACTGATGGAAACCAGCCATCGCCATCTGCTGCACGCGGAAGAAGGCACATGGCTGAATATCGACGGTTTCCATATGGGGATTGGTGGCGACGACTCCTGGAGCCCGTCAGTATCGGCGGAATTCCAGCTGAGCGCCGGTCGCTACCATTACCAGTTGGTCTGGTGTCAAAAAtaagtcgaccggctgctaacaaagcccgaaaggaagctgagttggctgctgccaccgctgagcaataactagcataaccccttggggcctctaaacgggtcttgaggggttttttgctgaaagccaattctgattagaaaaactcatcgagcatcaaatgaaactgcaatttattcatatcaggattatcaataccatatttttgaaaaagccgtttctgtaatgaaggagaaaactcaccgaggcagttccataggatggcaagatcctggtatcggtctgcgattccgactcgtccaacatcaatacaacctattaatttcccctcgtcaaaaataaggttatcaagtgagaaatcaccatgagtgacgactgaatccggtgagaatggcaaaagcttatgcatttctttccagacttgttcaacaggccagccattacgctcgtcatcaaaatcactcgcatcaaccaaaccgttattcattcgtgattgcgcctgagcgagacgaaatacgcgatcgctgttaaaaggacaattacaaacaggaatcgaatgcaaccggcgcaggaacactgccagcgcatcaacaatattttcacctgaatcaggatattcttctaatacctggaatgctgttttcccggggatcgcagtggtgagtaaccatgcatcatcaggagtacggataaaatgcttgatggtcggaagaggcataaattccgtcagccagtttagtctgaccatctcatctgtaacatcattggcaacgctacctttgccatgtttcagaaacaactctggcgcatcgggcttcccatacaatcgatagattgtcgcacctgattgcccgacattatcgcgagcccatttatacccatataaatcagcatccatgttggaatttaatcgcggcttcgagcaagacgtttcccgttgaatatggctcataacaccccttgtattactgtttatgtaagcagacagttttattgttcatgatgatatatttttatcttgtgcaatgtaacatcagagattttgagacacaacgtg | |

| P_T7 LacZ_ | Plasmid encoding LacZ under P_T7_ |
| --- | --- |
| P_T7_ Stability hairpin NonReg Terminator Kanamycin resistance cassette ColE1 origin    agatcaaaggatcttcttgagatcctttttttctgcgcgtaatctgctgcttgcaaacaaaaaaaccaccgctaccagcggtggtttgtttgccggatcaagagctaccaactctttttccgaaggtaactggcttcagcagagcgcagataccaaatactgttcttctagtgtagccgtagttaggccaccacttcaagaactctgtagcaccgcctacatacctcgctctgctaatcctgttaccagtggctgctgccagtggcgataagtcgtgtcttaccgggttggactcaagacgatagttaccggataaggcgcagcggtcgggctgaacggggggttcgtgcacacagcccagcttggagcgaacgacctacaccgaactgagatacctacagcgtgagctatgagaaagcgccacgcttcccgaagggagaaaggcggacaggtatccggtaagcggcagggtcggaacaggagagcgcacgagggagcttccagggggaaacgcctggtatctttatagtcctgtcgggtttcgccacctctgacttgagcgtcgatttttgtgatgctcgtcaggggggcggagcctatggaaaaacgccagcaacgcgatcccgcgaaattaatacgactcactatagggagaccacaacggtttccctctagaaataattttgtttaactttaaACCATGATTACGGATTCACTGGCCGTCGTTTTACAACGTCGTGACTGGGAAAACCCTGGCGTTACCCAACTTAATCGCCTTGCAGCACATCCCCCTTTCGCCAGCTGGCGTAATAGCGAAGAGGCCCGCACCGATCGCCCTTCCCAACAGTTGCGCAGCCTGAATGGCGAATGGCGCTTTGCCTGGTTTCCGGCACCAGAAGCGGTGCCGGAAAGCTGGCTGGAGTGCGATCTTCCTGAGGCCGATACTGTCGTCGTCCCCTCAAACTGGCAGATGCACGGTTACGATGCGCCCATCTACACCAACGTGACCTATCCCATTACGGTCAATCCGCCGTTTGTTCCCACGGAGAATCCGACGGGTTGTTACTCGCTCACATTTAATGTTGATGAAAGCTGGCTACAGGAAGGCCAGACGCGAATTATTTTTGATGGCGTTAACTCGGCGTTTCATCTGTGGTGCAACGGGCGCTGGGTCGGTTACGGCCAGGACAGTCGTTTGCCGTCTGAATTTGACCTGAGCGCATTTTTACGCGCCGGAGAAAACCGCCTCGCGGTGATGGTGCTGCGCTGGAGTGACGGCAGTTATCTGGAAGATCAGGATATGTGGCGGATGAGCGGCATTTTCCGTGACGTCTCGTTGCTGCATAAACCGACTACACAAATCAGCGATTTCCATGTTGCCACTCGCTTTAATGATGATTTCAGCCGCGCTGTACTGGAGGCTGAAGTTCAGATGTGCGGCGAGTTGCGTGACTACCTACGGGTAACAGTTTCTTTATGGCAGGGTGAAACGCAGGTCGCCAGCGGCACCGCGCCTTTCGGCGGTGAAATTATCGATGAGCGTGGTGGTTATGCCGATCGCGTCACACTACGTCTGAACGTCGAAAACCCGAAACTGTGGAGCGCCGAAATCCCGAATCTCTATCGTGCGGTGGTTGAACTGCACACCGCCGACGGCACGCTGATTGAAGCAGAAGCCTGCGATGTCGGTTTCCGCGAGGTGCGGATTGAAAATGGTCTGCTGCTGCTGAACGGCAAGCCGTTGCTGATTCGAGGCGTTAACCGTCACGAGCATCATCCTCTGCATGGTCAGGTCATGGATGAGCAGACGATGGTGCAGGATATCCTGCTGATGAAGCAGAACAACTTTAACGCCGTGCGCTGTTCGCATTATCCGAACCATCCGCTGTGGTACACGCTGTGCGACCGCTACGGCCTGTATGTGGTGGATGAAGCCAATATTGAAACCCACGGCATGGTGCCAATGAATCGTCTGACCGATGATCCGCGCTGGCTACCGGCGATGAGCGAACGCGTAACGCGAATGGTGCAGCGCGATCGTAATCACCCGAGTGTGATCATCTGGTCGCTGGGGAATGAATCAGGCCACGGCGCTAATCACGACGCGCTGTATCGCTGGATCAAATCTGTCGATCCTTCCCGCCCGGTGCAGTATGAAGGCGGCGGAGCCGACACCACGGCCACCGATATTATTTGCCCGATGTACGCGCGCGTGGATGAAGACCAGCCCTTCCCGGCTGTGCCGAAATGGTCCATCAAAAAATGGCTTTCGCTACCTGGAGAGACGCGCCCGCTGATCCTTTGCGAATACGCCCACGCGATGGGTAACAGTCTTGGCGGTTTCGCTAAATACTGGCAGGCGTTTCGTCAGTATCCCCGTTTACAGGGCGGCTTCGTCTGGGACTGGGTGGATCAGTCGCTGATTAAATATGATGAAAACGGCAACCCGTGGTCGGCTTACGGCGGTGATTTTGGCGATACGCCGAACGATCGCCAGTTCTGTATGAACGGTCTGGTCTTTGCCGACCGCACGCCGCATCCAGCGCTGACGGAAGCAAAACACCAGCAGCAGTTTTTCCAGTTCCGTTTATCCGGGCAAACCATCGAAGTGACCAGCGAATACCTGTTCCGTCATAGCGATAACGAGCTCCTGCACTGGATGGTGGCGCTGGATGGTAAGCCGCTGGCAAGCGGTGAAGTGCCTCTGGATGTCGCTCCACAAGGTAAACAGTTGATTGAACTGCCTGAACTACCGCAGCCGGAGAGCGCCGGGCAACTCTGGCTCACAGTACGCGTAGTGCAACCGAACGCGACCGCATGGTCAGAAGCCGGACACATCAGCGCCTGGCAGCAGTGGCGTCTGGCTGAAAACCTCAGCGTGACACTCCCCGCCGCGTCCCACGCCATCCCGCATCTGACCACCAGCGAAATGGATTTTTGCATCGAGCTGGGTAATAAGCGTTGGCAATTTAACCGCCAGTCAGGCTTTCTTTCACAGATGTGGATTGGCGATAAAAAACAACTGCTGACGCCGCTGCGCGATCAGTTCACCCGTGCACCGCTGGATAACGACATTGGCGTAAGTGAAGCGACCCGCATTGACCCTAACGCCTGGGTCGAACGCTGGAAGGCGGCGGGCCATTACCAGGCCGAAGCAGCGTTGTTGCAGTGCACGGCAGATACACTTGCTGATGCGGTGCTGATTACGACCGCTCACGCGTGGCAGCATCAGGGGAAAACCTTATTTATCAGCCGGAAAACCTACCGGATTGATGGTAGTGGTCAAATGGCGATTACCGTTGATGTTGAAGTGGCGAGCGATACACCGCATCCGGCGCGGATTGGCCTGAACTGCCAGCTGGCGCAGGTAGCAGAGCGGGTAAACTGGCTCGGATTAGGGCCGCAAGAAAACTATCCCGACCGCCTTACTGCCGCCTGTTTTGACCGCTGGGATCTGCCATTGTCAGACATGTATACCCCGTACGTCTTCCCGAGCGAAAACGGTCTGCGCTGCGGGACGCGCGAATTGAATTATGGCCCACACCAGTGGCGCGGCGACTTCCAGTTCAACATCAGCCGCTACAGTCAACAGCAACTGATGGAAACCAGCCATCGCCATCTGCTGCACGCGGAAGAAGGCACATGGCTGAATATCGACGGTTTCCATATGGGGATTGGTGGCGACGACTCCTGGAGCCCGTCAGTATCGGCGGAATTCCAGCTGAGCGCCGGTCGCTACCATTACCAGTTGGTCTGGTGTCAAAAAtaagtcgaccggctgctaacaaagcccgaaaggaagctgagttggctgctgccaccgctgagcaataactagcataaccccttggggcctctaaacgggtcttgaggggttttttgctgaaagccaattctgattagaaaaactcatcgagcatcaaatgaaactgcaatttattcatatcaggattatcaataccatatttttgaaaaagccgtttctgtaatgaaggagaaaactcaccgaggcagttccataggatggcaagatcctggtatcggtctgcgattccgactcgtccaacatcaatacaacctattaatttcccctcgtcaaaaataaggttatcaagtgagaaatcaccatgagtgacgactgaatccggtgagaatggcaaaagcttatgcatttctttccagacttgttcaacaggccagccattacgctcgtcatcaaaatcactcgcatcaaccaaaccgttattcattcgtgattgcgcctgagcgagacgaaatacgcgatcgctgttaaaaggacaattacaaacaggaatcgaatgcaaccggcgcaggaacactgccagcgcatcaacaatattttcacctgaatcaggatattcttctaatacctggaatgctgttttcccggggatcgcagtggtgagtaaccatgcatcatcaggagtacggataaaatgcttgatggtcggaagaggcataaattccgtcagccagtttagtctgaccatctcatctgtaacatcattggcaacgctacctttgccatgtttcagaaacaactctggcgcatcgggcttcccatacaatcgatagattgtcgcacctgattgcccgacattatcgcgagcccatttatacccatataaatcagcatccatgttggaatttaatcgcggcttcgagcaagacgtttcccgttgaatatggctcataacaccccttgtattactgtttatgtaagcagacagttttattgttcatgatgatatatttttatcttgtgcaatgtaacatcagagattttgagacacaacgtg | |
